# Mental compression of spatial sequences in human working memory using numerical and geometrical primitives

**DOI:** 10.1101/2020.01.16.908665

**Authors:** Fosca Al Roumi, Sébastien Marti, Liping Wang, Marie Amalric, Stanislas Dehaene

**Affiliations:** Cognitive Neuroimaging Unit, CEA, INSERM, Université Paris-Sud, Université Paris-Saclay, NeuroSpin center, 91191 Gif/Yvette, France; Collège de France, 11 Place Marcelin Berthelot, 75005 Paris, France; Institute of Neuroscience, Key Laboratory of Primate Neurobiology, Shanghai Institutes for Biological Sciences, Chinese Academy of Sciences, China; CAOS Lab, Department of Psychology, Carnegie Mellon University, 5000 Forbes Avenue, Pittsburgh, PA 15213

## Abstract

How does the human brain store sequences of spatial locations? The standard view is that each consecutive item occupies a distinct slot in working memory. Here, we formulate and test the alternative hypothesis that the human brain compresses the whole sequence using an abstract, language-like code that captures the numerical and geometrical regularities of the sequence at multiple nested levels. We exposed participants to spatial sequences of fixed length but variable regularity, and asked them to remember the sequence in order to detect deviants, while their brain activity was recorded using magneto-encephalography. Using multivariate decoders, each successive location could be decoded from brain signals, and upcoming locations were anticipated prior to their actual onset. Crucially, sequences with lower complexity, defined as the minimal description length provided by the formal language, and whose memory representation was therefore predicted to be more compressed, led to lower error rates and to increased anticipations. Furthermore, neural codes specific to the numerical and geometrical primitives of the postulated language could be detected, both in isolation and within the sequences. These results suggest that the human brain detects sequence regularities at multiple nested levels and uses them to compress long sequences in working memory.

## Introduction

Although non-human primates are able to learn sophisticated behavioral rules (Mansouri et al., 2020), the human species seems to be endowed with a deeper ability to discover the complex nested structures that underlie the information present in the environment (Dehaene et al., 2015). Such structural learning, indeed, is essential in order to acquire language, science, mathematics or music – activities which are all unique to humans. Even infants are able to quickly extract patterns from a stream of syllables, infer abstract regularities from a small number of examples, and generalize them to new items (Kabdebon et al., 2015; Marcus et al., 1999; Saffran, 2003; Xu and Tenenbaum, 2007).

A prominent hypothesis is that the capacity to represent embedded structures may be specific to the human species (Fitch, 2014; Hauser et al., 2002). Indeed, several comparative studies found that, while human and non-human primates exhibit similar performance in sequence processing whenever sequential relations, statistical properties or simple non-adjacent dependencies suffice to perform the task (Hauser et al., 2001; Milne et al., 2016, 2018; Newport et al., 2004; Ravignani et al., 2013; Sonnweber et al., 2015; Wilson et al., 2013, 2017), important differences are observed when the sequences involve embedding or recursion (Ferrigno et al., 2020; Fitch, 2004; Jiang et al., 2018; Wang et al., 2015). For instance, in a spatial motor task, although macaque monkeys could learn a supra-regular grammar and generalize it to novel sequences, they need a training period of thousands of trials to achieve the performance level that preschool children reach in only a few trials (Jiang et al., 2018). Thus, although the acquisition of simple embedded sequences may not be out of reach of macaque monkeys (Ferrigno et al., 2020; Jiang et al., 2018), the difference between human and non-human primates may originate, at least in part, in human’s ability to rapidly acquire complex sequential rules.

In the present research, which is part of series of behavioral and brain-imaging studies of spatial sequence learning (Amalric et al., 2017; Wang et al., 2019), we test the specific hypothesis that, when they learn a spatial sequence, human subjects make use of a language-like system of nested rules of variable complexity. To boost intuition, imagine that you have to view a sequence of 8 locations which are successively flashed on screen, in order to reproduce this sequence after a delay. This is the classical “Corsi blocks” neuropsychological test of spatial working memory, and the standard view is that each consecutive item should occupy one slot in working memory (Baddeley, 2003; Baddeley and Hitch, 1974; Botvinick and Watanabe, 2007; Hurlstone et al., 2014). Since the sequence length of 8 exceeds the typical human working-memory span, it would be difficult to remember it. Imagine, however, that you discover a strong regularity in the sequence -- perhaps the first four items form a square, and the next four draw another square. Mentally remembering this sequence as “2 squares”, thus compressing the information into an internal language-like expression, would considerably facilitate its memorization. In the present work, we test whether working memory is organized as a flat structure of slots or if it is structured as a language that allows to compress information.

In previous research (Amalric et al., 2017; Wang et al., 2019), we formalized the latter idea by proposing a hypothetical “language of thought” (Fodor, 1975) for sequences. The proposed language is akin to a mini computer language: it consists of primitives and rules whose combination can express any sequence of locations on an octagon (see figure 1). The central idea is that the successive locations are not just encoded by their spatial coordinates, independently of each other. Rather, the transitions between items are also encoded and if those transitions form a regular sequence, then those numerical regularities are also detected and encoded. The language thus comprises two types of instructions: geometrical primitives of rotation and symmetry (e.g. next item on the octagon, symmetrical item over the vertical axis, etc; figure 1C) as well as an operator of repetition, akin to the “for” loop in programming languages, that can repeat instructions a certain number of times, possibly with variations (see Supplementary Materials for a complete description). Since the language allows these instructions to be nested, it can express multiple levels of embedded repetitions and represent concepts such as “square”, “rectangle”, “two squares”, “zig-zag”, etc, through a combination of numerical and geometrical information.

**Figure 1:**
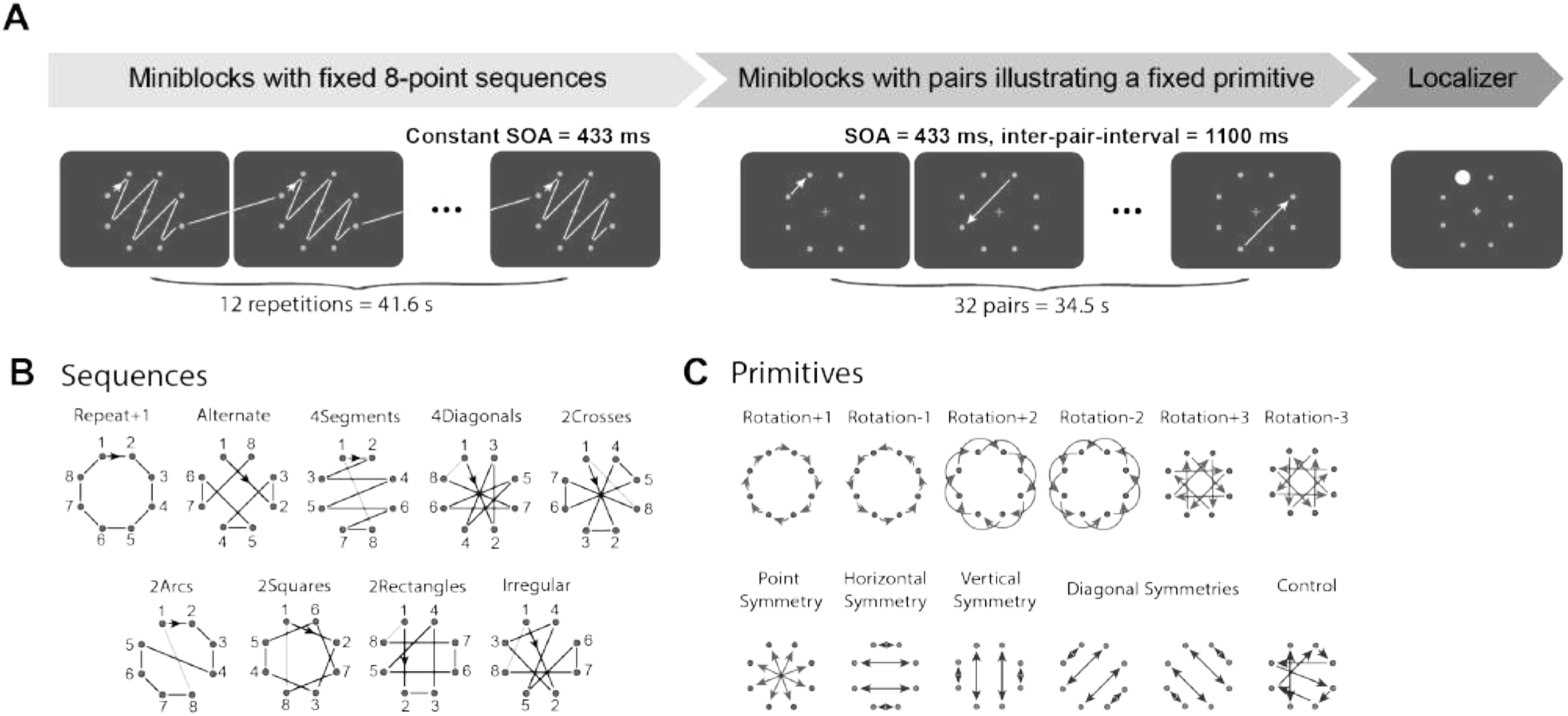
Experimental paradigm and stimuli. A. The experiment was divided into three parts: (1) In the sequence part, during each mini-block, a fixed spatial sequence of 8 locations was repeated 12 times; (2) In the primitive part, during each mini-block, 32 pairs of two successive locations illustrating a given primitive were presented; In both cases, participants were asked to report when they had identified the sequence or rule governing the pairs, and to click whenever they detected a violation. Finally (3), in the localizer block, dots were flashed at random locations on the octagon, and the data was used to train a location decoder. B. The nine 8-location sequence templates used in the sequence part. Presentation order is indicated by arrows. Actual sequences were generated by varying the starting point, rotation direction and/or symmetry axis. C. The pairs of locations illustrating each of the eleven primitive rules presented in the primitive part. Arrows indicate the first and the second element of each pair.

Our hypothesis is that, whenever they perceive a spatial sequence, human subjects attempt to represent it in memory by searching for the simplest mental program that can generate it. Thus, a simple prediction is that the difficulty of memorizing a sequence should not be proportional to its length, but to the length of the shortest mental program in the language of thought. Thus, sequences with a compact program should be easily memorized, even though their length may exceed the typical memory span of ~7 items, whereas at the other extreme, sequences with no compact description other than the mere list of locations would be harder or even impossible to learn. The underlying hypothesis, which has been previous proposed and tested in many other contexts (Chater and Vitányi, 2003; Feldman, 2000; Li and Vitányi, 1993; Mathy and Feldman, 2012; Romano et al., 2013), is that the brain operates as a “compressor” of incoming information that tries to select the minimal description for incoming stimuli, and that minimal description length is therefore a good predictor of psychological complexity. Hereafter, we refer to minimal description length as the “language-of-thought complexity” or LoT-complexity of a sequence.

In a first behavioral study, we evaluated the hypothesis that participants encode spatial sequences using the proposed language of geometry (Amalric et al., 2017). Participants were presented with sequences of 8 dots organized as a regular octagon and spanning a large range of complexities. They only saw the beginning of these sequences and had to predict the next locations. The results indicated that LoT-complexity was a good predictor of behavior. Specifically, the less complex the sequence was (as indicated by a greater compression rate), the more subjects were able to anticipate on the upcoming item, even on the first trial where they had never seen the entire sequence. Furthermore, error rate increased linearly as a function of minimal description length, and the error patterns across ordinal positions in the sequence was compatible with the nested structure of the expression postulated by the formal language. Another group (Yildirim and Jacobs, 2015) also showed how a similar compositional language for spatial sequences could account for the transfer of abstract sequence knowledge from the visual to the auditory modality. However, unlike the present work, they did not model participants’ sensitivity to geometrical regularities since the presented locations did not form regular geometrical shapes.

In a follow-up fMRI experiment on geometrical sequences (Wang et al., 2019), we merely asked participants to move their eyes to each item while the same 8-location sequences were presented on an octagon. Again, behavior indicated that gaze anticipation was inversely related to minimal description length and reflected each sequence’s nested structure. Activity in the dorsal part of inferior prefrontal cortex correlated with the amount of compression, while the right dorsolateral prefrontal cortex encoded the presence of embedded structures. Those brain regions belonged to a network that is distinct from but close to the areas involved in natural language processing, consistent with previous results indicating a dissociation of the neural circuits involved in mathematical thinking and in natural language processing (Amalric and Dehaene, 2017; Maruyama et al., 2012; Varley et al., 2005). Although the content of sequences could not be decoded from fMRI signals, multivariate pattern analyses provided indirect evidence that the activity patterns in dorsal prefrontal cortex became increasingly differentiated as the sequences were learnt.

fMRI is notoriously insensitive to the fine-grained timing of neural activity, and thus failed to directly probe the precise temporal unfolding of language-like rules that our theory predicted. In the current study, we aimed to further probe the existence of an abstract, language-like representation of geometrical sequences in the human brain by using magnetoencephalography (MEG), a sensitive technique with much higher temporal resolution than fMRI. We used time-resolved multivariate decoding (King and Dehaene, 2014) and representation similarity analysis (RSA) (Kriegeskorte et al., 2008) in order to examine if the postulated geometrical and numerical primitives could be decoded at the precise moment in the sequence where the postulated language of geometry suggests that they should be deployed. Thus, we exposed human participants to several repetitions of geometrical sequences of variable LoT-complexity. To ensure that they memorized the sequence, we asked them to click as soon as they had identified the repeated sequence, and to detect occasional sequence violations. We then tested if MEG signals were sensitive to the postulated numerical and geometrical primitives and to the proposed measures of sequence complexity and hierarchical structure. We identified markers of an anticipated representation of the sequence items and assessed their modulation by sequence LoT-complexity. Moreover, we exposed participants to multiple exemplars of each primitive operations in isolation, and we probed if those primitives could be extracted from the MEG signals. This approach allowed us to determine if the sequences were indeed represented as nested repetitions of primitive rules.

## Results

### Primitive rules: behavioral results

During the primitive part of the experiment, participants were exposed to a succession of two dots forming a pair illustrating a given hypothetical primitive operation (e.g. all pairs had a vertical symmetry). For instance, in a given block, for all pairs, the second dot was always vertically symmetrical to the first one, thus testing the postulate primitive of vertical symmetry. All 11 primitives of the language were tested (see the list in figure 1C, and the Methods section). We asked participants to click a button whenever they felt that they could predict the location of the second item in each pair. Those responses were converted into a measure of “encoding time” (figure 2A), expressed as the number of pairs that needed to be presented before the response occurred (if participants failed to respond, the maximum number of presentations, i.e. 32, was used). To assess if participants had understood the primitive rule and did not simply memorize the 8 pairs by rote, we also introduced a control condition in which the pairs were not driven by any general rule, and participants therefore had to memorize each of them. Participants performed very poorly in this condition, most of them failing to respond before the end of the run (i.e. after 32 pairs were presented). T-tests showed that, for all of the 11 proposed primitives, encoding time was shorter than for the control condition (all ps < 10^-5^). Such savings indicate that subjects detected all regularities.

**Figure 2:**
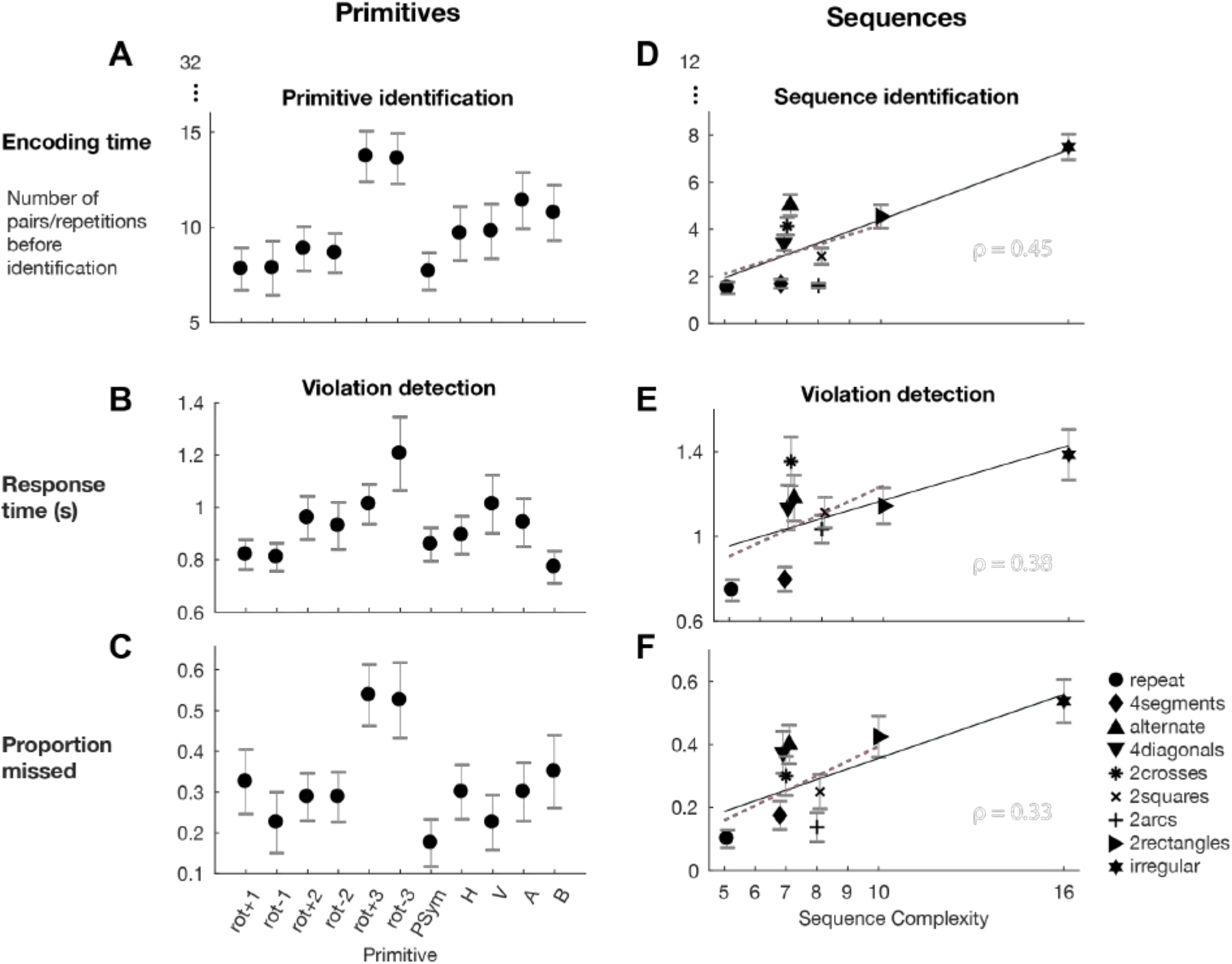
Behavioral performance and impact of sequence complexity. For the primitive part (left column) and the sequence part (right columns), graphs show (1) the mean encoding time, i.e. the mean number of repetitions that participants had seen before they reported identifying the rule or the sequence (A,D); (2) the performance in violation detection, as reflected in the mean response time to violations (top, B and E) and the proportion of missed trials (bottom, C and F). Error bars indicate one standard error of the mean (sem). For geometrical sequences, linear regression lines indicate the effect of theoretical sequence complexity. The red dotted lines were obtained when excluding the irregular sequence from the regression.

In the second half of each block, we introduced a violation detection task: participants were asked to press another button as fast as possible whenever they detected that the second dot of a pair was misplaced (figures 2B and C). In the control condition, participants missed 95% of those violations, whereas the average miss rate never exceeded 53% for the 11 genuine geometrical primitives. Again, all the differences relative to control were significant (all ps < 10^-4^).

Taken together, those results show that participants identified all of the primitive rules used to form the pairs and could use them to detect outliers. To determine if all rules were processed with the same ease, we ran repeated-measures ANOVA on three dependent behavioral measures: encoding time, violation detection time, and violation miss rate. This analysis revealed significant differences among the primitives (respectively F(10) =5.26, p < 10^-4^; F(10) = 3.31, p = 0.001; and F(10) = 4.00, p = p< 10^-4^). Tukey post-hoc tests on encoding time and miss rate indicated that the primitives of rotation ±3 were significantly more difficult than the counterclockwise rotation-1, the point symmetry and the vertical symmetry (see figure 2).

In summary, behavioral measures indicated that the participants could detect all of the postulated geometrical primitives, but that, contrary to our initial assumptions (Amalric et al., 2017) those primitives may not be strictly equivalent in complexity, with rotation±3 being more difficulty to detect (for a similar conclusion, see Romano et al., 2018a).

### Geometrical sequences: behavioral results

During the geometrical sequence block, subjects were repeatedly exposed to 8-item sequences. Two behavioral measures of sequence complexity were obtained. First, subjects were asked to press a button whenever they felt that they had identified the repeating sequence precisely enough to predict the next item. We again analyzed the encoding time, defined as the number of sequence repetitions that the participants needed before responding (if participants failed to respond, the maximum number of repetitions, i.e. 12, was used). As predicted, encoding time increased with our theoretical measure of sequence complexity (i.e. LoT-complexity), i.e. minimal description length in the proposed language (Spearman rank-correlation ρ = 0.45, t(19)=9.6, p < 10-7; Pearson correlation coefficient r = 0.70, t(19) = 20.7, p < 10-13; figure 2D; note that the results remained significant even when the most irregular sequence was excluded: Spearman rank-correlation ρ =0.22, t(19)= 3.1, p = 0.006; Pearson correlation coefficient r = 0.30, t(19)=4.3, p <10-3) (figure 2D).

After ten repetitions of a given sequence, i.e. during the last two presentations, a violation could occur (i.e. a single dot was misplaced, off by 3 locations on the octagon). Participants were asked to press a button as fast as possible when they detected it. The violation detection times again exhibited a positive correlation with LoT-complexity (figure 2E; Spearman rank-correlation ρ = 0.38, t(19)=7.1, p < 10^-5^; Pearson correlation coefficient r = 0.40, t(19)=6.7, p<10^-5^; when excluding the irregular sequence: Spearman rank-correlation ρ = 0.33, t(19)= 6.2, p <10^-5^; Pearson correlation coefficient r = 0.28, t(19)= 5.2, p <10^-4^). The effect was also detectable in error rates: as the sequence LoT-complexity increased, so did the number of missed violations (figure 2; ρ = 0.33, t(19)=6.3, p<10^-5^; r =0.43, t(19)=7.1, p < 10^-6^; excluding the irregular sequence, Spearman rank-correlation ρ = 0.18, t(19)= 2.4, p =0.03; Pearson correlation coefficient r = 0.28, t(19)= 3.8, p =0.001). Indeed, the effect on errors was very large: subjects missed less than 10% of deviants in the simplest sequence, but more than 50% of them in the most complex sequence, consistent with the idea that the latter exceeded their working memory span and they had trouble memorizing it in full. Since all sequences were of the same length (8 locations), those results cannot be explained by classical slot-based models of working memory, and argue in favor of the proposed language-of-thought hypothesis.

Overall, these results converge with the ones obtained using explicit predictions of the next item (Amalric et al., 2017) or using implicit eye-movement anticipations (Wang et al., 2019). They indicate that the more complex the sequences, the harder it is to predict the next locations, and therefore, the harder it is for participants to detect violations. Nevertheless, the correlations with minimal description length were modest, suggesting that our measure of sequence LoT-complexity, based on the proposed language, may not be ideal. Indeed, the behavioral results on the primitive part indicated that not all primitives should be considered as equally easy to encode. Specifically, rotation±3 was harder to process than the other primitives, and sequences involving this primitive could be harder to process. Furthermore, participant’s behavior may also be modulated by other parameters such as the spatial distance between items. To determine the contribution of these two variables to behavioral performance, we ran a stepwise regression (see *Methods*) to assess which linear model minimized the AIC (Akaike Information Criterion). The best model of encoding times and response times was one that included both LoT-complexity and presence of rotation±3. For the proportion of missed violations, the best model contained all three predictors: LoT-complexity, presence of rotation±3 and distance. Thus, the proposed language should be slightly amended to allow for a complexity that varies with distance and primitive type.

To obtain a more empirically driven measure of complexity, we computed the intercorrelation between our three dependent measures. Both detection time and missed rate were well predicted by encoding time, with correlations similar to those obtained from our theory-driven measure of LoT-complexity (respectively ρ=0.54, t(19)=10.5, p<10^-8^, r=0.51, t(19)=9.8, p<10^-8^, and ρ =0.44, t(19)=5.7, p<10^-4^, r =0.48, t(19)=5.4, p<10^-4^). We thus extracted the first principal component of those 3 dependent behavioral measures (figure 2D, E and F). Although determined in a purely data-driven manner, this empirical measure of complexity showed a robust correlation with the theory-driven LoT-complexity (ρ=0.56, r = 0.75). Deviations from a perfect line were due to deviations for the alternate, 4diagonals and 2crosses sequences, which could be explained by the fact that those sequences either contained the rotation±3 primitive or involved long spatial distances. In the rest of this paper, we shall use both theoretical and empirical complexity measures as predictors of brain activity, with the expectation that the empirical measure derived from behavior should be a better predictor.

### Decoding the successive locations of each sequence item

We first tested whether MEG signals contained decodable information about the successive locations of each sequence item. At each time point, based on the 306 sensor measures, we trained an estimator to decode the angular position of the presented item (flashed at time 0s). This position decoder was trained on data independent of the 8-item sequences and for which no anticipation could be formed (i.e. data from a localizer block, where locations were randomly intermixed, as well as data from the first item of each pair in the pair block, which was unpredictable; see *Methods* for details).

As shown in figure 3A, position decoding was at chance prior to stimulus presentation, but rose suddenly ~70 ms after stimulus onset, and peaked at 150 ms. The generalization-across-time matrix (figure 3B) revealed both a diagonal, indicating an unfolding sequence of stages, and a partial square pattern indicating a sustained maintenance of location information in brain signals (King and Dehaene, 2014). This location decoder, trained on independent data, was then applied to the 8-item sequence data. Successful generalization was observed: at each time step, the decoder successfully identified, with above-chance accuracy, the location of the current item in the sequence (figure 3C). To maximize the decodability of individual items in each sequence, we retrained a location decoder, but this time on the average brain responses in the time window of maximal decodability (from 100 to 200ms after the stimulus onset). We then tested it on each individual sequence. Figure 3D presents, for each sequence separately, the relative amount of times that the decoder predicted each location for each ordinal position. The pattern obtained from these predictions tightly paralleled the actual spatiotemporal profile of each sequence (second line of figure 3D), with only some added spatial uncertainty (i.e. spreading of the decoding to the two locations surrounding the correct one on the octagon). Control analyses showed that this decoding did not arise from residual eye movements, which were minimal (see Supplementary Materials).

**Figure 3:**
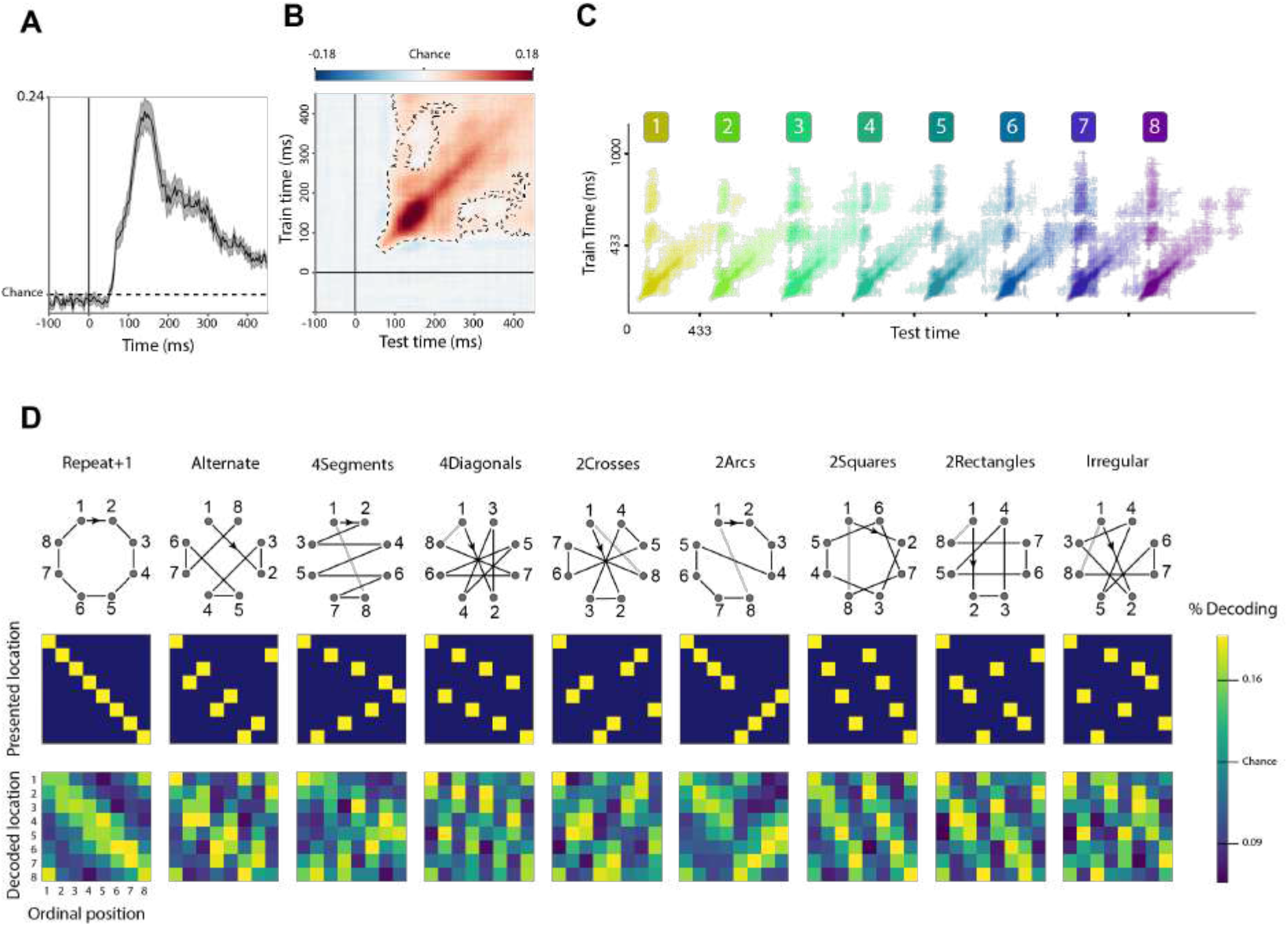
Decoding successive sequence locations from MEG signals. A. Performance in decoding stimulus location as a function of time following the flash of a dot at a given location. Maximal decoding performance is achieved at ~150 ms. B. Average generalization-across-time matrix showing the location decoding score as a function of training time (y axis) and testing time (x axis). The dashed lines indicate p < 0.05 cluster-level significance, corrected for multiple comparisons (see *Methods* for details on the cluster-based permutation analyses). C. Thresholded (p < 0.05, corrected) decoding matrix plot showing when each of the 8 successive sequence items could be decoded (averaged across subjects). D. Decoding of each of the nine 8-location sequences. For each sequence, the top matrix shows the stimuli at each ordinal position, and the bottom matrix shows the proportion of times a given spatial location was decoded.

Prior research on predictive coding has demonstrated that predictable stimuli elicit a reduced brain response, yet a more faithful representation as reflected by a higher decoding accuracy (Kok et al., 2012; Summerfield and de Lange, 2014). Here, we would expect this pattern to be present for easily predictable sequences with low LoT-complexity, but not for the more complex ones. To test this prediction, we examined if there was a linear modulation of the average decoding score as a function of LoT-complexity. We ran a linear regression on the decoding score as a function of theoretical and empirical LoT-complexity for each participant and determined if the regression coefficients associated to LoT-complexity were significantly negative, indicating that more complex sequences elicited a less precise internal representation of the successive locations. A small modulation was found as a function of empirical complexity (one tailed t-test t(19) = -2.0, p=0.0297) but not of LoT-complexity (one tailed t-test p = 0.1, t(19) = -1.3).

### Anticipation and its modulation by sequence structure

Our next goal was to determine if anticipatory information was present in MEG even prior to actual stimulus presentation, as previously demonstrated by others (Demarchi et al., 2019; Ekman et al., 2017; Kok et al., 2014, 2017). Indeed, the two behavioral tasks that participants had to perform during the experiment encouraged them to anticipate the location of the next stimuli in order to detect violations. The theory predicted that such anticipations would decrease with sequence LoT-complexity, indicating that more complex sequences were increasingly harder to predict.

To test for the presence of anticipation and its modulation by sequence LoT-complexity, we assessed the performance of the position decoder prior to the presentation of each sequence item (time 0ms). Importantly, since there was evidence of spillover of the decoding to nearby locations on the octagon, we controlled for distance to the previous item. To this aim, we defined an anticipation score as the difference in the decoding score at the upcoming location and at the symmetrical but non-stimulated location, which was equidistant from the previous location (see *Methods* and figure 4A). This anticipation score was computed for all sequence items (figure 4B) then averaged across the training time window 100-200 ms (figure 4C) which corresponds to the maximal performance of the position decoder (figure 3A). We ran a cluster-based permutation test in the temporal window between the presentation of the preceding item and the anticipated item (i.e. from -430ms to 0ms). The anticipation score was significantly positive during two time-windows, from -220 to -115 ms and from -85 to 0 ms prior to stimulus presentation. Thus, participants did memorize and anticipate sequences, at least in part, based on a location-specific code. Additional analyses, presented as supplementary materials, excluded a contribution of eye movements to those anticipation signals (see figure S1). Although a contribution of micro-saccades below the spatial resolution of our eye tracker cannot be fully excluded, note that the latter would still constitute evidence for anticipation.

**Figure 4:**
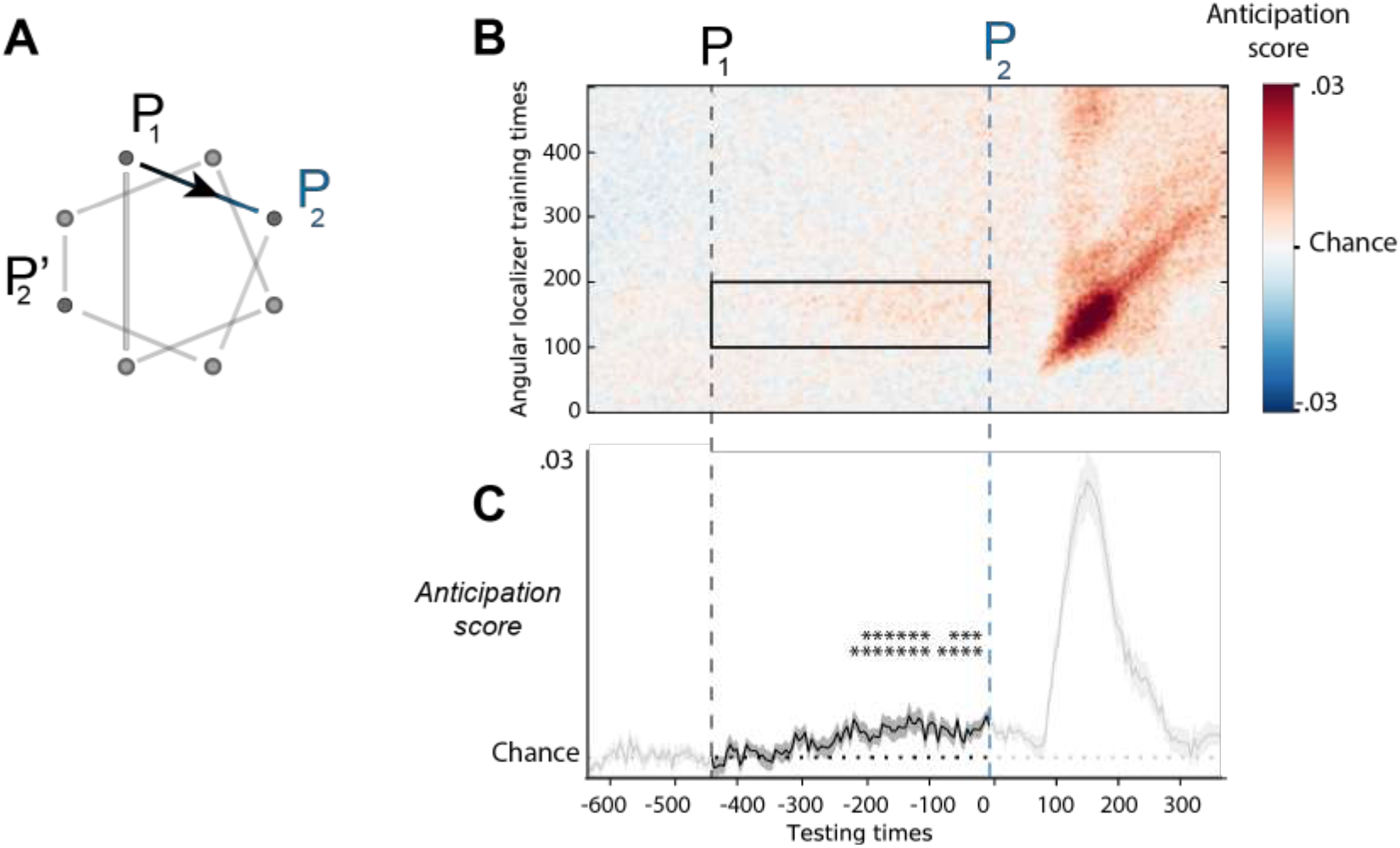
Detecting an anticipation of sequence locations from MEG signals. For each training time in the range 0-500 ms, we tested whether the location decoder could detect the stimulus location in a time window ranging from 650 ms before to 400 ms after stimulus presentation. A. To control for distance from the previous stimulus location (P1), an anticipation score was computed by measuring the decoding at the correct stimulus location (P2) and subtracting the decoding at the equidistant, symmetrical, non-stimulated location (P’2). B. Generalization-across-time matrix, indicating that a decoder trained in the window 100-200 ms after P2 shows a small amount of generalization prior to stimulus presentation. C. Temporal evolution of the anticipation score, obtained by averaging the anticipation score over training times ranging from 100 to 200 ms (selected region from panel B). The anticipation score was significantly above chance starting ~250 ms before stimulus presentation, indicating that the brain anticipates P2 even before it appears.

We then used the anticipation score to probe the internal representation of the sequences. If predictive mechanisms are modulated by sequence structure, then the anticipation score should be increasingly smaller as the sequence gets more complex. We computed the average anticipation score for each sequence (except 4diagonals and 2crosses; see *Methods*), and ran a linear regression as a function of complexity. We observed a significant decrease of anticipation score with LoT-complexity (one-tailed t-test t(19)=-2.1, p=0.025) and empirical complexity (one-tailed t-test t(19)=-2.0, p=0.028). This result provides indirect evidence for the hierarchical representation postulated by the language of geometry, as it shows that expectation mechanisms, measured by the anticipation score, are modulated by the overall LoT-complexity of the sequence. Note that this effect did not depend on distance, but solely on LoT-complexity, as a similar regression showed that the anticipation score did not vary significantly with the distance between the anticipated item and the preceding one (t(19)=-1.4, p>0.1).

Previous research has shown that brain activity in language areas is modulated by the nested structure of language, such that activity varies sharply at the boundary of sentence constituents such as noun phrases (Nelson et al., 2017). We thus wondered if a similar effect occurred with the language of geometry. To do so, we therefore compared the anticipation scores of the items that, according to our postulated language, open a constituent, to the ones that are inside a component (i.e. inside a series of repeated instructions). The analysis was run only on the 4segments and 2squares sequences. The Repeat1, alternate and irregular sequences were excluded from it as their postulated representation does not involve a nested syntactic structure. We also excluded the 2arcs sequences since all constituent opening corresponded to distance-4 transitions (see *Methods*). Finally, we also discarded the complex 2rectangle sequence, since its anticipation score was not significantly different from zero. Perhaps due to this reduced data set, and the ensuing lack of statistical power, this analysis did not reveal any significant difference as a function of syntactic structure.

### Decoding geometrical operations

The language-of-geometry hypothesis predicts that participants encode spatial sequences not only in terms of the item’s specific location, but also in terms of high-level geometrical primitives such as symmetries and rotations. To directly probe the representation of the sequences in terms of the language of geometry, we attempted to decode, from the brain signals, the elementary geometrical primitives postulated by our formal language.

To do so, we first examined if the 11 primitive operations could be decoded when they were presented in isolation. Using the trials that illustrated each primitive (see figure 1C), we trained a decoder to determine, based on the brain signals, which of the 11 primitive operations was presented (*see Methods*). Our hypothesis was that participants actively apply the operation (e.g. symmetry around the vertical axis) after the presentation of the first element of the pair, in order to predict the location of the second. To characterize the temporal dynamics of the primitive operation code, the primitive decoder was trained and tested at different time-points. We cross-validated across runs to exclude decoder overfitting due to temporal proximity, and we ran a sliding window over the epochs to increase the signal-to-noise ratio (*see Methods*). Significance was assessed by a cluster-based permutation test on the window -200 to 600 ms. As shown in figure 5A, performance was significantly above chance (p < 0.05 cluster-level significance) for an extended time window which peaked ~200-300 ms following the presentation of the first element of the pair, but actually started ~50 ms before that presentation, suggesting that the block structure enabled participants to anticipate on the forthcoming geometrical transformation. Thus, those results indicate that human brain signals contain decodable information about the type of geometrical transformation that links a location *n* and the next one *n*+1 in a sequence, and do so, in a familiar block or sequence context, even before the second sequence item is presented.

**Figure 5:**
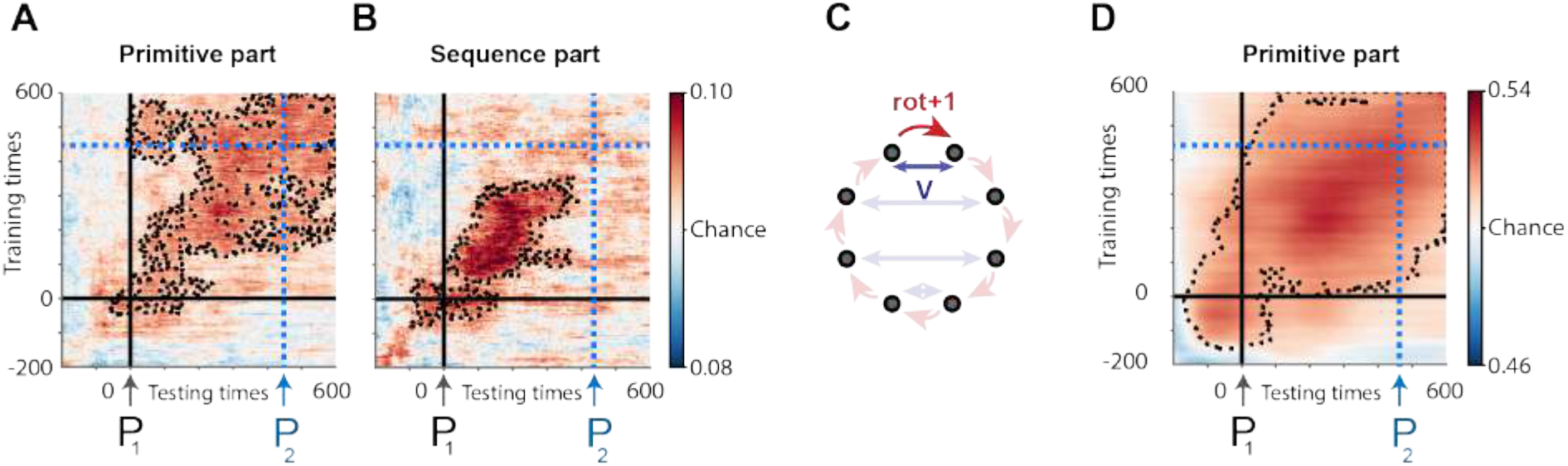
Decoding the geometrical primitives. Decoders were trained to decode which of the 11 postulated geometrical transformations was applied at a given time between two consecutive locations P1 and P2 (the onset of the first location is marked by t = 0 ms). A, B: Average generalization-across-time matrices showing the decoding score as a function of training time (y axis) and testing time (x axis) for the 11 primitive operations in the primitive part of the experiment (A) and in the sequence part (B). Panel C illustrates how trials with the same start and end locations may be classified as a rotation or a symmetry depending on the context in which they are presented. Panel D shows the performance of a binary decoder for rotation versus symmetry on such trials with identical start and end locations, taken from the primitive part of the experiment. The dashed lines indicate p < 0.05 cluster-level significance over the -200 ms – 600 ms time-window, corrected for multiple comparisons (see *Methods*).

The 11 geometrical primitives in Figure 1C are all perfectly balanced in terms of starting point and endpoint, and their decoding is therefore unconfounded by retinotopic stimulation. However, by definition, they involve different *pairs* of locations, and some (e.g. +1, +2, +3) differ in terms of the distance between the two elements of the pair. It is unclear whether this represents a genuine artifact, because the time-window when they were decoded preceded the presentation of the second item of the pair, hence came before it could have any physical effect (e.g. perceived motion). Still, perhaps what we decoded was the distance of the attentional movement involved to anticipate the next location, rather than the abstract geometrical transformation itself. We therefore wondered whether, in the extreme case, geometrical transformations could be decoded even the pair of starting points and endpoints were strictly identical. We capitalized on the fact that the same pairs of dots could appear in the context of either a rotation block or a symmetry block. For instance, a dot moving from the top left to the top right location can be construed as either a rotation around the octagon (+1), or as a symmetry with respect to the vertical axis (see figure 5C for illustration). We selected all trials corresponding to such pairs and asked whether, for equal starting and ending locations, the brain signals still contained decodable information about their putative internal encoding as a rotation or as a symmetry (*see Methods*, figure 5C). Figure 5D shows that this was indeed the case. These decoding results suggest that, over and above any location or distance information, the neural patterns associated with abstract geometrical operations of rotation and symmetry can be disentangled when primitive operations are considered in isolation.

We then determined whether, based on the description provided by the postulated formal language, the same primitive operations could be decoded in the context of a sequence. We extracted 800 ms epochs centered on specific locations in the sequence, and labelled them by the subsequent primitive operation involved at the lowest level of the sequence description (i.e. the one whose application predicted the next location). We successfully trained a decoder for the 11 primitives, as indicated by an above-chance generalization-across-time matrix (figure 5B). However, when we replicated the above training and testing on a balanced set of rotation and symmetry pairs, we could not detect, at above-chance level, a code separating the rotation and symmetry operations – perhaps because of the smaller number of trials involved.

We then tried to determine if the primitive code generalized from the primitive part of the experiment, when primitive operations are presented in isolation, to the sequence part of the experiment, when elementary primitive operations are recruited in the context of a program. In order to avoid any bias due to a particular block or run, we trained this decoder on the micro-averaged trials over the 4 runs (see *Methods*) and tested it on the sequence data. The generalization-across-time matrices did not exhibit any significantly above chance clusters, suggesting that the neural code for the primitives when they are presented in isolation is not directly replicated in the context of a full sequence, but is modified in the sequence context (see General Discussion).

In summary, the MEG decoding analysis provided evidence for an abstract encoding of rotations and symmetries, independently of the visual features of the stimuli, both in the primitive and in the sequence parts of the experiment. However, this code did not generalize from the primitive to the sequence part.

### Characterizing the code for primitive operations with RSA

To supplement the decoding approach, we adopted a Representation Similarity Analysis approach (Kriegeskorte et al., 2008). Relative to the decoding approach, RSA presents the potential advantage of using all trials, while carefully modeling and therefore disentangling the superimposed influences of visual and higher-level factors on brain activity. We therefore used RSA to further test the existence of a neural code for abstract geometrical primitives, independently of a location-specific code. In both the primitive and the sequence parts of the experiment, we selected epochs ranging from -500 ms to 1000 ms around a selected pair of locations, and for which that transition was encoded by an unambiguous primitive according to our proposed language. We then computed the full representational dissimilarity matrix between epochs, separately for each primitive operation and each location on screen (*see Methods*). These empirical dissimilarity matrices were then regressed as a function of several theoretical predictors (see figure 6). One predictor captured the part of the experiment to which the stimuli belonged (primitive or sequence; here called “block type”). The specific geometrical primitive, as well as the primitive type (rotation or symmetry), were also modeled, both within and across experimental parts. Additional predictors were introduced to capture retinotopic or visuospatial factors (the locations of the first and the second item of the pair, and the distance between them). Consistently with our initial decoding results, we observed a strong contribution of those visual predictors to MEG signals. The code for item spatial location had a similar temporal dynamics and latency as the location decoding performance (compare figure 6 and figure 3A). The distance between two consecutive spatial locations also had a significant influence on brain signals at later time points, peaking after the position of the second item. The regression coefficients for block type peaked in the period preceding the presentation of the first item, consistent with the differences in the temporal structure of the trials belonging to the two types of blocks.

**Figure 6:**
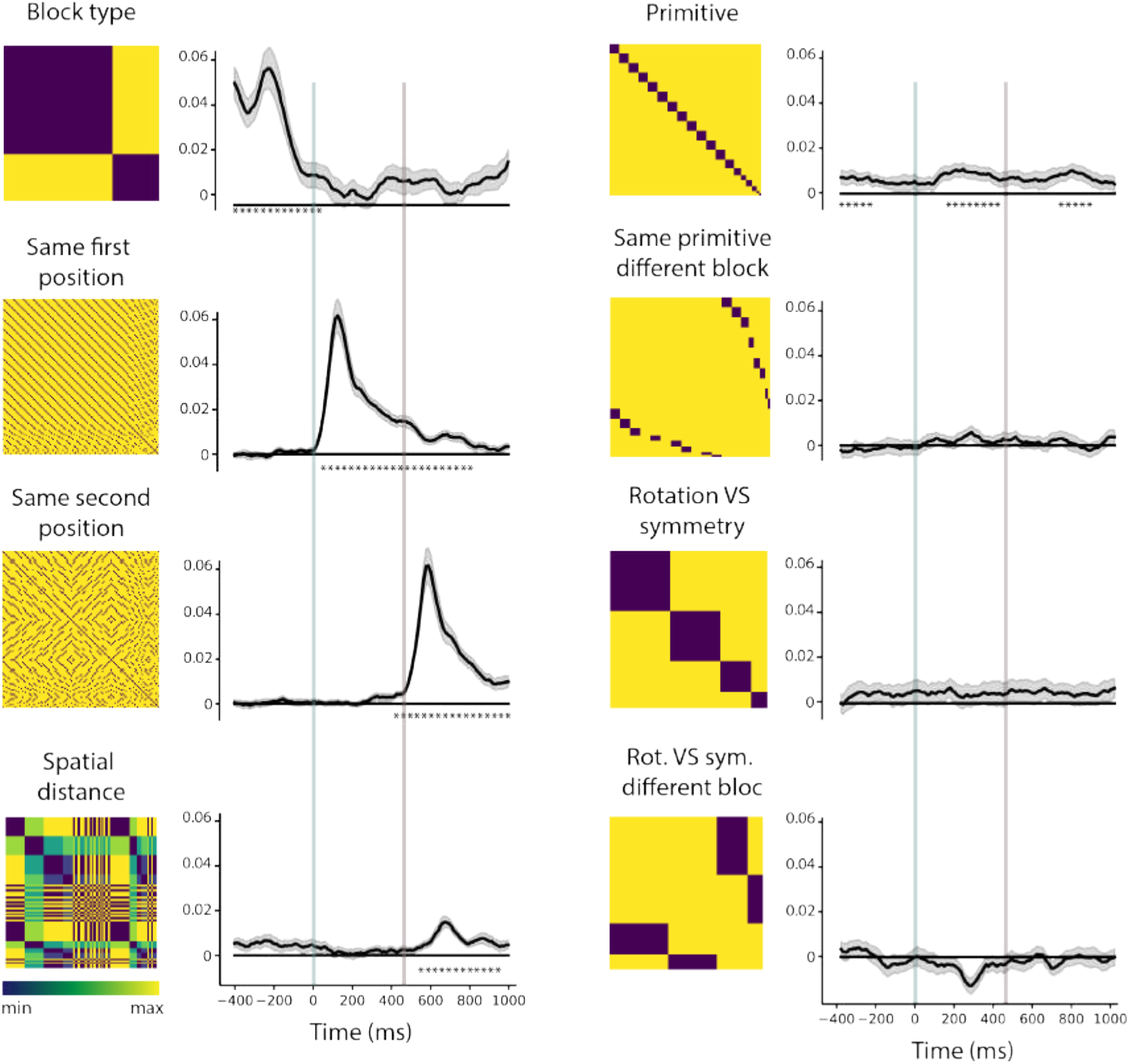
Disentangling visual and higher-level codes for spatial sequences using RSA. In this representational similarity analysis (RSA), a linear combination of multiple theoretical dissimilarity matrices was used to model the empirical dissimilarity matrices obtained at different times surrounding a pair of locations, thus investigating the temporal dynamics of sequence coding. The left column represents the theoretical dissimilarity matrices. In the matrix, epochs are sorted by block type (primitive or sequence part of the experiment), then by primitive operation, and finally by the presented pair of locations on screen. Each empirical dissimilarity matrix, at a given time *t*, is regressed as a function of these theoretical predictors and the graphs at right column represent the corresponding regression coefficients. Stars denote significant time windows based on cluster-based permutation tests. The vertical lines denote the onset of the first (blue) and second (grey) items of a pair.

Most importantly, a cluster-based permutation test on the 0–1 sec time window confirmed that a representation of abstract geometrical primitives also influenced brain activity. Regression coefficients for the primitive operation were significantly positive in the window between 100 ms and 400 ms, i.e. in between the presentation of the two items of the pair, during which an application of the corresponding transformation allowed to predict the location of the second item. This representation was reactivated after the second item was presented (significant cluster from 710 to 870 ms).

We ran a separate cluster-based permutation test from -400 ms to 0 ms in order to test for anticipation effects. Indeed, we found a significant cluster on the window -400 ms to -250 ms suggesting that the primitive operation was represented before the first item of the pair was presented. We then tried to determine if the neural code for primitive operation was similar across experimental parts. Although a mild peak was observed (figure 6), it did not reach significance after correction for the time interval tested. Thus, consistent with the decoding results, the neural representation of geometrical primitives differs in the context of a sequence and when presented in isolation. Finally, in this analysis, no significant similarity between primitive operations belonging to the same type (rotation or symmetry) could be found.

### Decoding ordinal position in subsequences

According to the proposed hypothesis, the internal representation of the sequence is not limited to a mere list of locations, or even of transitions between locations, but takes the form of a language-like code that expresses nested structures such as “2 squares”, or more precisely “2 groups of 4 items, each linked by a +2 operation”. Our language predicts that, when remembering such a sequence, an internal numerical code, akin to a “for i=1:n loop”, must unfold in the participants’ brain, keeping track of how many times a given geometrical transformation has been applied. Thus, we investigated the presence, in human MEG signals, of a neural code for ordinal position within a subsequence of items, ranging from 1 to a maximum of 4 in our sequences; such a code has been previously observed in a simpler context in both human and non-human primates (Kutter et al., 2018; Nieder, 2012; Nieder et al., 2006) and has been postulated in some models of working memory (Botvinick and Plaut, 2006; Botvinick and Watanabe, 2007). We refer to those analyses as “ordinal position decoding”, although we readily acknowledge that, since our sequence unfolded at a fixed pace, the coding of number and elapsed time could not be distinguished. However, crucially, since each sequence of 8 locations continually repeated itself without gaps, the presence of such an ordinal code cannot simply reflect elapsed time since the beginning of a block. Rather, finding such a code would provide support for the postulated parsing of the sequence into multiple subgroups of locations linked by a common geometrical transformations (similar to the parsing of language sequences into nested phrases; see Ding et al., 2016).

Our analyses focused on the 2squares and 2arcs sequences, which according to our proposal contain 2 groups of 4 items; and on the 4segments and 4diagonals sequences, which contain 4 groups of 2 items. In each case, we trained and tested a decoder on sequence data labeled by ordinal position at the lowest level of the postulated mental expression (respectively a four-way decoder for ordinal positions 1-4; and a binary decoder for ordinal positions 1 versus 2). In both cases, we trained and tested our decoders at different times, thus obtaining a full generalization-over-time matrix, while cross-validating across runs. The results showed above-chance decoding around the diagonal (figure 7A and C, left column), suggesting that ordinal position was indeed encoded in MEG signals. Decoding performance was above chance before the stimulus was actually presented, compatible with the fact that the sequences repeated continuously and could therefore be anticipated.

**Figure 7:**
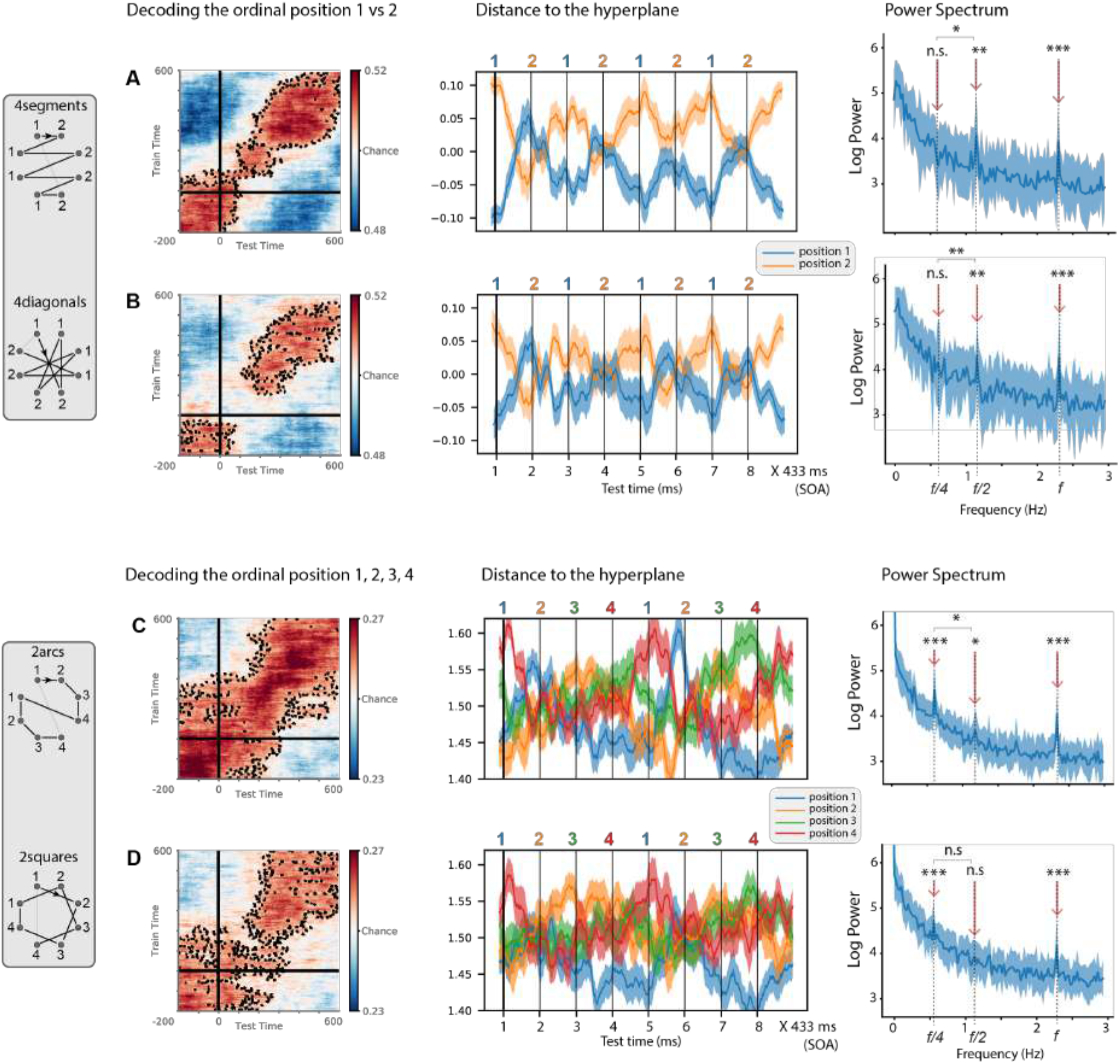
Decoding ordinal position within a sequence component. Decoders were trained to predict the ordinal positions 1-2 in the inner groups of sequences 4segments and 4diagonals (panels A-B), and ordinal positions 1-4 in the inner groups of sequences 2arcs and 2squares (panels C-D). Panels A and C were obtained by cross-validating across blocks and B and D by generalizing across sequences (e.g. from 4segments to 4diagonals or vice-versa). Left column, mean generalization-across-time (GAT) performance for the ordinal position decoder (accuracy score): each cell of this GAT matrix indicates generalization of a decoder trained on data at a given training time (y axis) and tested on data from a given testing time (x axis). 0ms corresponds to item onset. Middle column, time course of the average output of ordinal decoders trained in a time window 300-500 ms, during a sequence of 8 consecutive items. Although noisy, those curves show a decoding peak ~400 ms after the corresponding ordinal item, and a clear rhythmicity every 2 items for the top sequences (A-B), and every 4 items for the bottom sequences (C-D). The y axis shows the distance to the ordinal classification hyperplane for each decoder. The onsets of the sequence items are indicated by vertical lines, and times are reported from the onset of the first item (thick vertical line). Blue, orange, green and red lines indicate 1^st^, 2^nd^, 3^rd^ and 4^th^ positions. Right column, power spectrum obtained from the Fourier transform of those time courses throughout the full presentation blocks for the two sequences. Statistics are provided at frequencies f = 2.31 Hz, f/2=1.15 Hz and f/4 = 0.58 Hz.

We then tested the prediction that the internal code for ordinal position is abstract and therefore identical regardless of the particular geometrical primitive that is called for at that position. In other words, we asked whether the code for a sequence is “factored out” into separate codes for ordinal position and for the particular geometrical primitive applied at this position, as predicted by our language model and as previously shown for non-spatial visual sequences by Liu et al., 2019. If the codes for the ordinal positions 1-4 were similar across, say, the 2squares and 2arcs sequences, indicating a progression through the four successive stages of the internal loops respectively drawing a square (+2 operation) and an arc (+1 operation), then we should be able to train an ordinal position decoder on one sequence and generalize it to the other. Indeed, we found above-chance generalization across time and across sequences (figure 7B and D, left column), indicating that the code for ordinal position is, at least in part, independent of the specific geometrical transformation involved.

To characterize the temporal variations of this code during the entire sequence presentation, we applied the ordinal position decoders to MEG data from left-out runs that were not used for training. As an estimator of classifier performance, rather than the percentage of correct classifications, we computed a more sensitive estimate, the signed distance to the classification hyperplane for each predicted ordinal position. Since the decoding score was above chance for training and testing times from 300 ms to 500 ms, we averaged the predicted distances on this time window. The results (middle column) indeed showed oscillations compatible with a sequential unfolding of an ordinal position code across the entire sequence, either going 1-2-1-2-1-2-1-2 for sequences 4segments and 4 diagonals, or 1-2-3-4-1-2-3-4 for sequences 2squares and 2arcs. This was true for decoders trained on both sequences (figures 7A and 7C, middle column) and also when generalizing across sequences (figures 7B and 7D, middle column). To test if the ordinal code indeed oscillated at the appropriate frequency, we computed the log power spectrum of the distance time-series for each condition. We then determined if there was a significant peak at the component frequency (2arcs and 2squares, f/4 = 0.58 Hz; 4diagonal and 4segments, f/2 = 1.15 Hz) by comparing the log power at that frequency to the neighboring frequencies (*see Methods*). In the case of 2squares and 2arcs, the test was performed on the averaged log power over the 4 ordinal position decoders. In every condition, the test was significant at the predicted frequency, indicating the presence of groups of 2 or 4 items depending on the sequence (p<0.01). Furthermore, a 2 x 2 ANOVA of the log power difference, with factors of frequency (f/2 or f/4) and sequence type (components of size 2 vs 4), showed a significant interaction (p < 0.01), both when training on both sequences and when generalizing across sequences, indicating that the power was significantly stronger at the expected than at the inappropriate frequency (see *Methods*).

Instead of ordinal number, the brain might have solely encoded the position of the first and the last item in each group. However, we could reject this hypothesis because, as shown in Figure S2, significant oscillations were also present, with a peak at the appropriate time, for the intermediate positions 2 and 3 in 4segment and 4diagonal sequences. Nevertheless, examination of the cross-generalization between the ordinal codes for groups of 2 and for groups of 4 items suggested that the internal code for sequences also included first versus last information (see Figure S3). First, the ordinal code for the first and the last component positions of 2arcs and 2squares generalized to the first and the last sequence items of the 4segments and 4diagonals sequences. The power spectrum of the distance time-series showed a significant peak at f/2 which originates from the reactivation of the codes for the 1^st^, 2^nd^ and the 4^th^ (i.e. last) position (p<0.01 at f/2 for these positions). Second, the ordinal codes for 1^st^ and 2^nd^ position, extracted from 4segments and 4diagonals sequences, reactivated consistently for the 1^st^ and 2^nd^ ordinal positions of the 2arcs and 2squares sequences (as confirmed by a significant peak at f/4) but interestingly, the decoder for the 2^nd^ position also maintained a sustained activation, not only at the 2^nd^ position of 2arcs and 2squares, but also at their 3^rd^ and especially their 4^th^ position (Figure S3, bottom). Taken together, these results suggest that the ordinal decoding was partially driven by two different kinds of codes, one for ordinal number (1-2 or 1-2-3-4 depending on group size) and another for first versus last (which coincided with ordinal positions 2 or 4 depending on group size).

As a final control, to verify whether those sophisticated decoders would achieve above-chance performance on any kind of data, we trained and tested the very same decoders on data for which our language model predicts that there should be no subgroups of items, i.e. the repeat and irregular sequences. As predicted, the decoders now failed to identify a 1-2-1-2 structure (figure S4, panels A and B; and Supplementary materials). Interestingly, there was modest evidence for a 1-2-3-4 code only in the repeat sequence, suggesting that even when structure is absent (the items keep going around the octagon without any break), participants may encode them in memory as groups of 2 or 4 (perhaps as top versus bottom ones, left versus right, or according to the sides of the octagon). Crucially, the decoders found no evidence of ordinal coding in the irregular sequence.

## Discussion

The goal of the present study was to probe the internal representations that humans use to encode geometrical sequences of varying regularity. The simplest models of working memory for serial order assume either that sequences are encoded as simple associative chains linking one item to the next (e.g. TODAM model, Lewandowsky and Murdock Jr., 1989) or by storing the successive items in distinct memory slots, based on their position in the sequence (Botvinick and Watanabe, 2007; for review, see Hurlstone et al., 2014). If this was the case, however, all of our sequences would be encoded in a similar manner, as they all have the same length and only differ in the order in which the same eight locations are presented. Instead, we found evidence that participants mentally compress the sequences using their geometrical regularity, and have a better memory for those that can be compressed down to a lighter memory load. Indeed, both behavioral and brain-derived measures were modulated by LoT-complexity, as provided by the postulated formal language. Furthermore, using time-resolved decoding and RSA techniques, we identified three distinct types of codes: for the item’s specific spatial location, for the geometrical operation that characterizes the transition from one location to the next, and for their ordinal position within a group of items. We now discuss those points in turn, starting with evidence for those elementary primitives, and continuing with evidence that those can be combined into language-like expressions.

First, using multivariate decoding, we found that each of the successive retinotopic locations in a spatial sequence could be decoded (Figure 3). However, we also found that the brain did not stop at encoding specific locations, but also coded for the transitions between consecutive locations, in such a way that we could classify them into 11 abstract geometrical primitives (e.g. horizontal symmetry; rotation by 1 item counterclockwise around the octagon, etc) and decode them (Figure 5). Decoding of geometrical primitives worked both when considering primitive operations in isolation and in the context of a sequence. In addition to evidence from decoding, representational similarity analysis (RSA) allowed to factor out the contributions of retinotopic and abstract geometrical codes, and to demonstrate that, even if the first one dominates in the MEG signal, the second, although weaker, is significant (Figure 6).

The results thus indicate that the human brain uses multiple redundant codes for sequences: although each sequence could have been stored as a series of 8 retinotopic locations, subjects also encoded it as a series of geometrical transformations. Our findings supports the existence of a mental repertoire of abstract geometrical concepts and its automatic deployment when a sequence must be encoded in memory. It fits with previous behavioral evidence that even young children and adults with limited access to formal schooling in mathematics already use these geometrical primitives when memorizing sequences (Amalric et al., 2017). All humans, starting at an early age, may have access to “core knowledge” of basic geometrical concepts, comprising multiple basic capacities for spatial navigation, shape recognition and topology (Dehaene et al., 2006; Spelke et al., 2010).

Another prediction of our formal language is that the human brain uses such geometrical regularities to segment the sequences: if a series of successive locations can be encoded by the repeated application of the same transformation, (e.g. three consecutive applications of the +2 operation to draw a square on the octagon), then subjects compress it internally using an internal repetition operator, akin to a “for” loop in programming languages. As a result, we predicted that, although the sequences unfolded continuously and without any break, we should be able to identify a mental code for the successive ordinal positions 1,2,3… within such a loop. In agreement with this hypothesis, MEG signals contained decodable information about ordinal position within a subgroup of locations (Figure 7). The ordinal code was activated in a periodic manner (figure 7 E-H) in agreement with the proposed rhythmicity of the sequence representation, which varied across sequences (groups of 2 or 4). Such ordinal knowledge was not present in every sequence, but could only be decoded from the brain signals when the code was indeed predicted by our language model (compare figure 7 and control figure S4).

Future experiments could use variable sequence length, tempo and grouping in order to better specify the nature of the neural code involved. We consider most plausible that the subjects encoded each item’s ordinal number, between 1 and 4, because several previous experiments have revealed that such a numerical code is present in several cortical regions of the monkey (Nieder, 2012; Nieder et al., 2006; Ninokura et al., 2004) and human brain (Fias et al., 2007; Kutter et al., 2018; Nieder and Dehaene, 2009). However, given a fixed tempo, it is possible that we also decoded the elapsed time since the beginning of a group. Furthermore, we also found suggestive evidence for the decoding of the first and the last ordinal positions (which correspond to the opening and closing of a component, and may therefore reflect the distinct mental operations involved). This finding fits with the ubiquity of primacy and recency effects in working memory (Anderson et al., 1998; Hurlstone et al., 2014; Orlov et al., 2000; Terrace et al., 2003). Nevertheless, the ordinal code for intermediate positions 2 and 3 could also be decoded and recurred in a periodic manner (figure S4), indicating that the ordinal code was not limited to first versus last.

Importantly, all such codes point to the same conclusion, namely that the human brain does not stick to a flat, superficial representation of the sequence, but parses it into subsequences based on geometrical cues. In this respect, our results extend the conclusions obtained for linguistic structures in connected speech (Ding et al., 2016) to the domain of visuospatial sequences.

Another important postulate of our proposed language of thought is that different sequences can be encoded via a recombination of the same ordinal and geometrical codes. In a recent MEG study, Liu et al. (2019a) proposed the term “factorized codes” for such a situation in which a sequence is encoded as the combination of an abstract structural code (here for the ordinal number 1-4) and a specific content (here the specific geometrical transformation, e.g. +1 to generate an arc or +2 to draw a square). In their study, Liu et al. exposed participants to sequences of pictures and asked them to reorder them according to a learned rule. They showed that human MEG signals contain three different codes for item identity, item ordinal position in the sequence, and the ultimate target sequence after the requested mental transformation. These codes were partially separated in time and the hippocampus replayed the learnt structure, supporting the hypothesis of a factorized hippocampal representation of abstract structural sequence knowledge. Liu et al. suggested that abstract structure is encoded in the medial entorhinal cortex, while specific sensory representations are encoded in the lateral entorhinal cortex (Manns and Eichenbaum, 2006a; Whittington et al., 2019a). The current results are consistent with Liu et al’s (2019a) findings, because we found that the structural code that tracks ordinal position generalizes across sequences that make use of different geometrical primitives. This result suggests that ordinal knowledge is encoded in an abstract manner, independently and separately from the code used to represent the particular geometrical primitive. This subdivision of labor fits with our formal language, in which the same “for” loop, with an identical numerical pointer, can be applied to different geometrical primitives. The distinction between number and geometrical primitives suggests that the brain encodes sequences using factorized codes for those two different dimensions of sequence knowledge.

A final prediction of our language is that those primitives can be nested to form complex, recursive mental programs, and that working memory load is ultimately determined by the minimal description length of that program, once it is encoded in the proposed language-of-thought – a measure we called language-of-thought complexity (LoT-complexity). We obtained behavioral as well as MEG evidence in support of this prediction. Behaviorally, participants’ subjective feeling of remembering the sequence, as well as their objective capacity to detect an occasional location deviant, were correlated with LoT-complexity (figure 2). In MEG, we found that covert visual anticipation of the sequence items was modulated by sequence complexity, suggesting that expectation mechanisms can be more efficiently deployed when the sequence has a low LoT-complexity. The observation of anticipation signals fits within the predictive-coding framework, which proposes that the brain constantly tries to predict its sensory inputs (Bastos et al., 2012; Chao et al., 2018; Friston et al., 2003) and projects stimulus-specific templates in advance of the stimulus itself, as previously found for instance during associative learning tasks (Demarchi et al., 2019; Kok et al., 2014, 2017; Sakai and Miyashita, 1991) or during sequences of spatial locations with no structure (Ekman et al., 2017). Our own previous behavioral studies showed that human participants could anticipate the geometrical sequence items, both by explicitly pointing and by implicitly moving their gaze to their spatial location before they were actually presented, and that such anticipation varied according to the hierarchical structure of the sequences (Amalric et al., 2017; Wang et al., 2019). Wang et al. (2019) further showed that fMRI signals in bilateral inferior frontal gyrus (IFG) and dorsolateral prefrontal cortex (dlPFC) were modulated by LoT-complexity and by the actual amount of anticipation of the nested rules. These regions were previously shown to be engaged in the parsing of structured spatial sequences (Bor et al., 2003; Desrochers et al., 2015), but those studies considered only two levels of sequence regularity (structured *vs* unstructured), as opposed to the multiple levels of regularity and nesting studied here. The results from Wang et al. (2019) bolster the proposition of a hierarchical caudal-rostral organization of the prefrontal cortex to represent rules of increasing abstraction (Badre, 2008; Badre and D’esposito, 2009; Badre and Nee, 2018; Badre et al., 2010; Koechlin and Jubault, 2006; Koechlin et al., 2003). These prefrontal regions may encode the internal model of geometrical sequences and send top-down signals in order to pre-activate circuits in posterior cortices, generating the observed anticipations.

Several limits of the present work must be acknowledged. First, the formal language of geometry that we proposed provided only an imperfect, though significant, fit to the observed behavioral data (figure 2). Indeed, an empirical measure of complexity, derived from participants’ behavioral performance, provided a better predictor of MEG anticipatory activity. This deviation of behavior from theory could arise from the fact that, for simplicity, we assigned to all primitives the same weight in the computation of LoT-complexity. In reality, some primitives, such as rotation±3, which involve greater spatial distances, seemed to be empirically more difficult to process than others. Indeed, when adding a distinct weight for rotation±3 in a linear model, we obtained a better predictor of the behavioral data. In an augmented version of the theory, weights could be assigned to each primitive separately before computing sequence complexity (as proposed by Romano et al., 2018a).

A second limit is, although we provided indirect support for the proposed language through the effect of LoT-complexity on brain and behavior, we did not provide direct decoding evidence that several numerical and geometrical primitives could be jointly encoded in a complex program (e.g. 2 squares = 2 groups of 4 items, each linked by a +2 operation, separated from each other by a +1 operation). So far, we only managed to decode the lowest level of this hypothetical nested code, i.e. the transitions between consecutive items and their ordinal position within a local group of locations. Whether and how the brain “binds” the same representation at multiple nested levels to create a nested or “syntactic” code is a difficult question which has been debated at the theoretical level for over 30 years (e.g. Elman, 1990; Marcus et al., 1999; Smolensky and Legendre, 2006) and remains to be empirically resolved. In this context, it is noteworthy that the decoder of geometrical primitives, once trained on the data from the primitive part (where only pairs of locations were presented), did not generalize to the sequence part where the same primitives were embedded into longer sequences. This result was confirmed by RSA. Future research should disentangle two alternative explanations. First, this negative finding could be due to peripheral factors such as the different timing of the two parts (figure 1), or the possibility of head position changes across the experiment. However, it may also hint at a principled difference in the neural encoding of the same geometrical primitives when presented in isolation and in a sequence context. Indeed, a major difference is that, in the primitive part, a single primitive rule is considered at the time, for an entire block, whereas in the sequence part, the primitives have to be bound together with other primitives and with numerical information in order to form more complex expressions such as “2 squares”. Some theories of the neural codes for syntax postulate that, to embed an object inside a syntactic structure, the brain applies a tensor product operation between the neural codes for the object and for its role inside the structure (Smolensky, 1990; Smolensky and Legendre, 2006). This radical transformation would re-map the original neural vectors for primitives onto a very different direction in high-dimensional neural space, thus explaining why they can no longer be decoded by the original decoder once they are embedded in a sequence. Higher resolution recordings, possibly at the single-cell level, may be required to test this hypothesis.

Alternative neural codes may also have contributed to the observed difference between the pair and sequence parts of our experiment. Contrary to primitive pairs, sequences were repeated identically multiple times, thus facilitating memorization and perhaps fostering the use of other forms of sequence representation, over and above the proposed minimal set of rotations and symmetries. For instance, subjects may have encoded sequences using visual aids. Even though vertices or edges were never explicitly displayed, lines could have been mentally constructed linking the separate sequence items, and further integrated to form shape percepts (e.g. a mental image of a square) that may be encoded in early visual cortices (Gilbert and Li, 2013; Kok et al., 2014; Li et al., 2006; Roelfsema and Houtkamp, 2011). The complex shapes drawn by the full sequences may in turn be decomposed into elementary shapes such as squares, circles, triangles or rectangles (Leyton, 2001). Successive locations might also have been encoded as consecutive landmarks of trajectories travelled in an allocentric perspective. To memorize these trajectories, participants may build a cognitive map analogous to the one involved in egocentric spatial navigation (Aronov et al., 2017; Constantinescu et al., 2016; Hafting et al., 2005; O’Keefe and Nadel, 1978; Tolman, 1948).

## Conclusion

The present work fits with a long line of research, going back at least to the 19^th^ century (Aronov et al., 2017; Constantinescu et al., 2016; Hafting et al., 2005; O’Keefe and Nadel, 1978; Tolman, 1948), focusing on the “language of thought” hypothesis (Fodor, 1975), according to which humans encode complex concepts as nested combinations of a finite set of elementary primitives. This idea was initially supported by behavioral studies of human rule learning (Shepard et al., 1961). In concept learning experiments, the speed and efficiency of learning was shown to be modulated by the Boolean complexity of the rule, i.e. the length of its shortest logical expression as a formula with elementary logical operators and parentheses (Feldman, 2000). Working memory for sequences of digits was also found to be modulated by the possibility of reorganizing it into simpler chunks (Mathy and Feldman, 2012). Such research led to the general proposal that the human brain acts as a compressor of information in all sorts of domains, and always attempts to select the shortest expression that accounts for what it perceives (Chater and Vitányi, 2003; Feldman, 2000, 2003; Li and Vitányi, 1993; Romano et al., 2013). The present research adds further evidence in favor of this framework. Furthermore, very recently, the same formal language of geometry that we tested here was also found to successfully capture the regularities in binary auditory and visual sequences made of only two arbitrary sounds or pictures (Planton et al., 2020). Future research should examine three key open questions. First, to what extent can the same set of recursive language-like rules capture very different domains where humans excel, such as mathematics, music or language? Second, how are such rules implemented at the neural level? And third, are such codes uniquely developed in human, as postulated by some researchers (Dehaene et al., 2015; Fitch, 2014; Hauser et al., 2002), or can they also be observed in non-human primates?

## Acknowledgements

We acknowledge help from all the NeuroSpin support teams, and particularly Leila Azizi and Virginie van Wassenhove. We are grateful to Maxime Maheu, Pedro Pinheiro-Chagas and Darinka Trübutschek for their valuable comments. This research was supported by INSERM, CEA, Collège de France, the Bettencourt-Schueller Foundation and an ERC grant “Neurosyntax” to SD. It was performed on a platform of the France Life Imaging network, partly funded by Agence Nationale de la Recherche grant *ANR-11-INBS-0006*, and supported by the Bettencourt-Schueller Foundation and the Leducq Foundation.

## Methods

### Participants

20 participants (9 men; *M*_age_ = 24.6 years, *SD*_age_ = 3.7 years) with normal vision were included in the MEG experiment. In compliance with institutional guidelines, all subjects gave written informed consent prior to enrollment and received 90€ as compensation.

### Experimental protocol

#### General structure of the experiment

The main task, completed in the MEG Elekta acquisition device, was subdivided into 3 parts. To avoid biasing subjects towards specific primitives, the sequence part was performed first. The second part was dedicated to each of the primitive operations, and the third part was a localizer task with unpredictable locations, designed to train a decoder for spatial locations. During the 3 parts of the MEG experiment, white dots were flashed for 100ms on the vertices of an octagon while the subject was fixating a cross at the center of the screen. The MEG experiment was preceded by a short training (c.a. 20 minutes) to the geometrical sequences.

#### Training on sequences

As initial training, outside the MEG, the geometrical sequences were presented with a slower pace than the rest of the experiment: a stimulus onset asynchrony (SOA) of 700ms between consecutive dots, and a dot duration of 200ms. Each sequence was repeated until the participants pressed the space bar to report that they had memorized it. They were then asked to type in the 8 locations that followed the last item displayed on the screen. If they succeeded, the word ‘Bravo’ (congratulations) was presented on the screen and the next sequence started. If they failed, the word ‘Erreur’ (error) was displayed and the same sequence restarted. Participants were instructed that the same sequences would be presented to them during the main experiment. No training was provided on the primitive part of the experiment. The training part was meant to select participants able to quickly encode the geometrical sequences. Only the ones that had finished the training in less than 20 minutes were qualified for the main MEG experiment. 5 out of 25 participants did not manage to do the training part of the experiment in less than 20 minutes.

### MEG task

#### Geometrical sequences

The first part of the experiment was devoted to the geometrical sequences and was composed of 4 runs. 9 sequences (figure 1B) were composed of 8 non-repeating locations. The last 3, called ‘Memory sequences’, which were composed of 1, 2 or 4 spatial locations, were not analyzed in this study. During one run, each sequence of 8 locations was presented 12 times consecutively. The sequences presented in figure 1B were mere templates: to generate the actual sequences, the starting point and global direction of rotation were varied and balanced across runs. 4segment sequences were selected such that each of the 4 symmetry axes (horizontal, vertical and diagonal) would appear once. Participants had to perform two tasks. First, they had to report with a button press when they had identified the sequence and felt able to predict the next locations. Second, during the 11^th^ or the 12^th^ repetition, an item appeared at an unexpected location, and subjects had to report this violation with another button press as fast as possible. This task was added to maintain participants attention during the full block. MEG epochs containing such a violation were excluded from all analyses.

#### Primitive part

The second part of the experiment was devoted to the primitive operations. It was composed of 4 runs, subdivided in 12 mini-blocks for each of the 12 conditions. The 11 first ones followed elementary ‘primitive’ rules (figure 1C), because a simple geometrical operation allowed to determine the spatial location of the second item of a pair when given the first. The 12^th^ condition was a control condition in which no minimal rule allowed to do so, and participants could only memorize the 8 unrelated pairs in order to perform the task. A mini-block was composed of 32 pairs with a SOA of 433ms between the items of the pair and an inter-pair-interval of 1100ms. Each of the 8 pairs appearing 4 times in the mini-block.

The task was similar to the sequence part. Participants reported with a button press when they had identified the rule that allowed them to predict the location of the second item given the first. In addition, they had to detect as fast as possible when the second item did not appear at the expected location. Violations could only occur during the presentation of the last 8 items of the mini-block. Again, MEG epochs containing such a violation were excluded from all analyses.

#### Localizer part

The last part of the experiment was meant to train a spatial decoder for spatial position of the presented items. To do so, dots were flashed pseudo-randomly on the vertices of the octagon with SOA 433ms. Occasionally (1/20 dots on average) the color of the dot changed. The subject had to click as fast as possible to report this.

### MEG acquisition and preprocessing

#### MEG recordings

Participants performed the tasks while sitting inside an electromagnetically shielded room. The magnetic component of their brain activity was recorded with a 306-channel, whole-head MEG by Elekta Neuromag^®^ (Helsinki, Finland). 102 triplets, each comprising one magnetometer and two orthogonal planar gradiometers composed the MEG helmet. The brain signals were acquired at a sampling rate of 1000 Hz with a hardware highpass filter at 0.03Hz. The data was then decimated by a factor 4.

Eye movements and heartbeats were monitored with vertical and horizontal electrooculograms (EOGs) and electrocardiograms (ECGs). Subjects’ head position inside the helmet was measured at the beginning of each run with an isotrack Polhemus Inc. system from the location of four coils placed over frontal and mastoïdian skull areas.

#### Data cleaning: Maxfiltering

Bad MEG channels were identified visually in the raw signal and were provided to the MaxFilter software (ElektaNeuromag^®^, Helsinki, Finland) to compensate for head movements between experimental blocks by realigning all data to an average head position and to apply the signal space separation algorithm (Taulu et al., 2004) to suppress magnetic interference from outside the sensor helmet and interpolate bad channels.

#### Data cleaning: ICA

The rest of the analysis was performed with MNE Python (Gramfort et al., 2013; Jas et al., 2018). Oculomotor and cardiac artefacts were removed performing an independent component analysis (ICA). The components that correlated the most with the EOG and ECG signals were automatically detected. We then visually inspected their topography and correlation to the ECG and EOG time series to confirm their rejection from the MEG data.

#### Multivariate Pattern Analysis

The goal of the Multivariate Pattern Analysis (MVPA) decoding analyses was to predict from single-trial brain activity (*X*) a specific categorical (e.g. primitive identity) or continuous (e.g. angular position) variable (*y*) that represents the neuronal state corresponding to the participant’s mental representation. These analyses were performed following King et al’s preprocessing pipeline (King and Dehaene, 2014) implemented in MNE-python version 0.16 (Gramfort et al., 2013). Prior to model fitting, each channel at each time-point was z-scored across trials. Each estimator was fitted on each participant separately, across all MEG sensors using the parameters set to their default values provided by the Scikit-Learn package (Pedregosa et al., 2011). When the estimator was trained and tested on two different conditions, the whole training and testing sets were used to respectively fit and test the estimator. By constrast, when the decoder was trained and tested on non-independent data, we used either a 5-folds stratified cross-validation procedure to prevent overfitting or we cross-validated across block number. The reported scores are the average across crossvalidation folds.

#### Decoding of angular position

The spatial angular decoders were built from two ridge regressions used to decode the angular position Θ. One predicted sin(Θ) and the other cos(Θ). The angular decoding score was obtained by first computing the mean absolute difference between the predicted angle (Θ_pred_) and the true angle (Θ_true_). We subtracted to this score π/2 to obtain a score in the range of - π/2 and π/2 (chance = 0) (King and Dehaene, 2014). When we wanted to determine which of the 8 locations was predicted by the angular decoder, we binned its output into 8 evenly spaced angular bins.

To access the temporal organization of the neural representations, we computed the temporal generalization matrices. These matrices represent the decoding score of an estimator trained at time t (training time on the vertical axis) and tested with data from another time t’ (testing time on the horizontal axis).

#### Anticipation score

The anticipation score was defined as the subtraction of the number of times the correct position was predicted in advance, and the number of times the other position at the same distance from the preceding item was predicted (figure S1). This measure was designed to overcome a potential confound that comes from the fact that successive sequence items tend to be close to each other (average angle between two sequence items is 73°, s.d. = 28°). As the angular decoder output spreads on neighboring angles, it may predict above chance the spatial position of the next item. By construction, the anticipation score is immune to such a distance-based confound. The anticipation score was not defined when the next item is at distance 4 from the previous one. As this represents 50% of the transitions for 4diagonals and 2crosses sequences, we excluded these sequences from the linear regression anticipation analyses that we ran across participants.

#### Decoding of ordinal position in sequence component

We ran the ordinal decoder on smoothed data: we averaged the brain signals over a sliding window of 100 ms for every time-step (i.e. each 4 ms). We used Support Vector Machine classifiers to decode the ordinal position at the lowest level of the hierarchical code predicted by our language model (i.e. 1-2-1-2… or 1-2-3-4 depending on the sequence, see Figure 7). The decoders’ performance was estimated by computing their mean accuracy, averaged over participants.

#### Decoding of primitive identity

We used Support Vector Machine classifiers to decode which geometrical primitive was involved at a given transition between two locations, as predicted by our language model. Brain activity was smoothed as for the decoding of the ordinal position. When controlling for visual confounds, rotation±2 primitive was excluded from the trial set as no symmetries involved a distance of 2. Moreover, when only two categories were considered (e.g. ‘rotation’ and ‘symmetry’), the score of the decoder was provided in terms of the area under the curve. When decoding the 11 primitives, we used One-VS-rest multiclass classifiers and report their mean accuracy.

#### Representational similarity analysis

Representational similarity analysis (RSA) characterizes a neural representation by the similarity between the neural patterns elicited by a set of stimuli. To test a given model, we compare the representational similarity it predicts to the one measured from brain signals. In this experiment, we hypothesized that brain activity would reflect a superposition of visuospatial factors (item location, distance between consecutive items) and high-level geometrical primitives. To compute the similarity between two conditions, we smoothed the data over sliding windows of 100ms with 10ms steps. Epochs belonging to the even and odd run numbers were averaged separately to form two sets of evoked activities. The empirical similarity was determined as the Spearman rank correlation between the evoked activities of these two sets. The empirical dissimilarity (1-Spearman rank correlation) was then regressed as a function of the theoretical representational dissimilarities (predictor matrices were z-scored).

#### Fourier analysis and statistical assessment of the presence of a peak

To determine the periodicity of the activation of the ordinal code, we ran a fast Fourier transform on the distance to the hyperplane time-series using *scipy*’s *fftpack* module and computed the difference between the log-power at the test frequency and the log-power at neighboring frequencies. The log-power at f (2.31 Hz), f/2 (1.15 Hz) and f/4 (0.577 Hz) was compared to the average log-power for the 4 neighboring frequencies (frequency bin width: 0.024 Hz) using a student t-test. The difference in log-power for f/2 and f/4 was compared using a student t-test. To determine if there was an interaction between the amplitude at f/2 and f/4 and the component size, we used a linear mixed model (Matlab^®^ function *fitlme*) to fit the power difference as a function of component size, frequency and an interaction between frequency and component size. Subject number was considered as a random factor. We then ran an ANOVA for this linear mixed model (Matlab^®^ function *anova*).

#### Statistical analyses

All statistics reported in the text refer to group-level analyses. Tukey post-hoc tests were performed using the R software packages *nlme* and *multcomp*. We used multiple comparison corrections to assess multivariate decoding performance for the two-dimensional time by time generalization score matrices and for the simple decoding score time-courses. We considered temporal clusters and non-parametric one-sample *t*-tests estimated on 4096 permutations (Maris and Oostenveld, 2007) implemented by the *permutation_cluster_1samp*_test available in *mne.stats* package. A cluster was defined by adjacent time points. The cluster-level statistic was the sum of the sample-specific *t*-statistics that belonged to a given cluster. The alpha level of the sample-specific test statistic and of the cluster-specific test statistic were 0.05. For decoding performance curves, ‘*’ and ‘**’ indicate that resulting p-values are respectively < 0.05 and < 0.01. Dashed contours on temporal generalization correspond to p-value < 0.05 resulting from the permutation test. The student t-tests and the linear regressions were performed with Matlab^®^. Stepwise linear regressions were computed with stepwiselm Matlab^®^ function using an AIC criterion starting from a model with intercept.

## Supplementary materials

### Minimal description and hierarchical structure of the geometrical sequences

The proposed language of geometry includes instructions for repeating operations and for more complex repetitions with variations. For example, the sequence [+0, [+1]^7^] starts from the point 0 and repeats 7 times the operation rotation+1, thus describing a circle around the octagon. The sequence [[+0, H]^4^{-1}] builds a horizontal segment by applying the horizontal symmetry to 0, then iterates it four times, each time shifting the starting point according to rotation-1 – thus describing a zigzag or 4-segment sequence. The sequence [[+0, P]^4^〈+1〉] builds a diagonal from the point symmetry P connecting position 0 to 4, and then iterates it four times, shifting the two elements by rotation+1 to build each of the consecutive segments forming a star-shaped or 4-diagonals sequence. Further information on the language of geometry is provided in the supplementary material of Amalric et al. (2017). Table S1 provides a minimal description of each of the 9 sequences under study. The * sign indicates an ambiguity, because several descriptions with the same amount of compression exist for the considered sequence, depending on starting point, rotation direction and/or symmetry axis. Only the transitions with unambiguous geometrical or ordinal primitives were used for decoding.

**Table S1:**
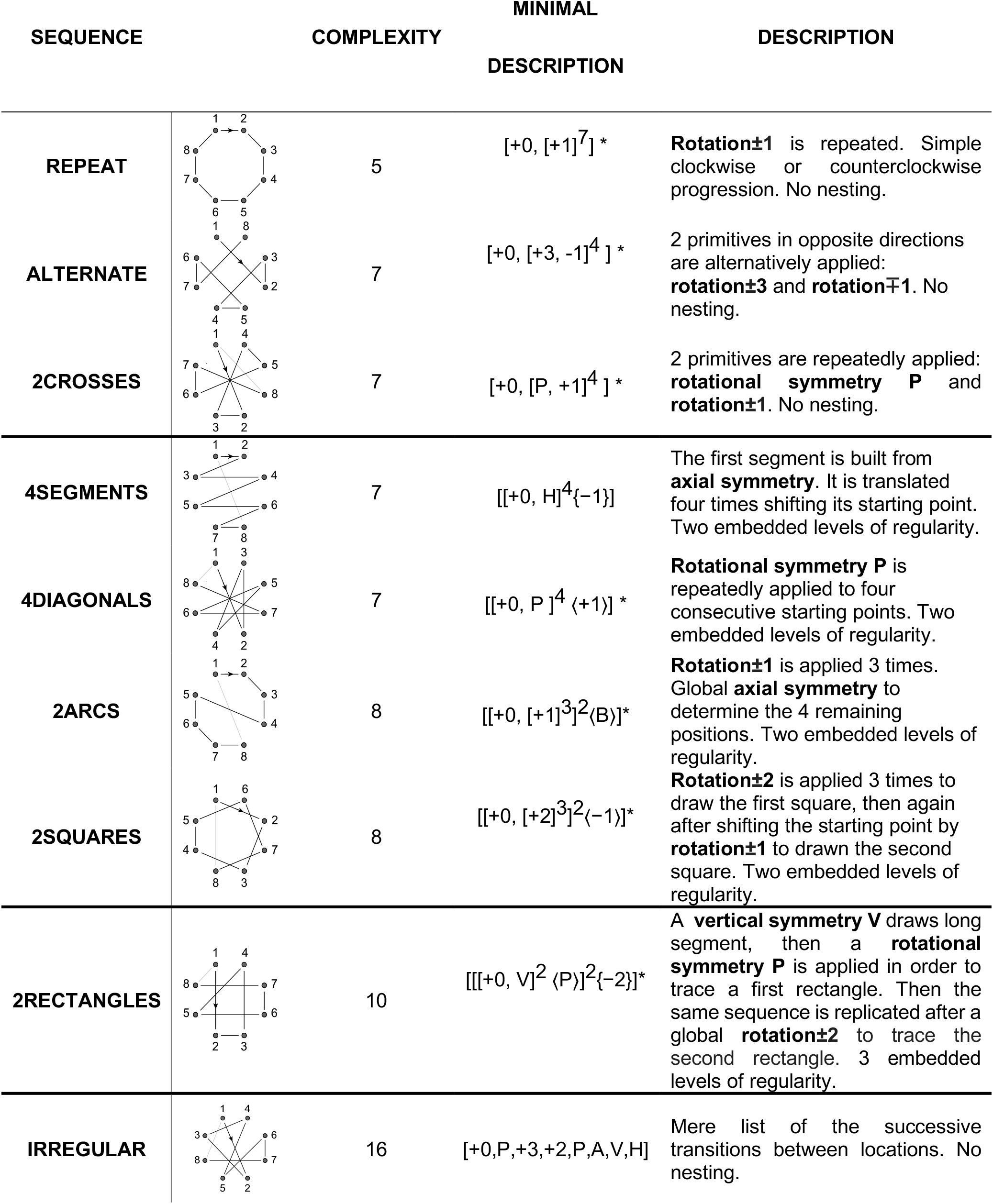
Description of the geometrical sequences

| SEQUENCE | COMPLEXITY | MINIMAL DESCRIPTION | DESCRIPTION |
| --- | --- | --- | --- |
| REPEAT | 5 | $[+0, [+1]^7]^*$ | **Rotation±1** is repeated. Simple clockwise or counterclockwise progression. No nesting. |
| ALTERNATE | 7 | $[+0, [+3, -1]^4]^*$ | 2 primitives in opposite directions are alternatively applied: **rotation±3** and **rotation∓1**. No nesting. |
| 2CROSSES | 7 | $[+0, [P, +1]^4]^*$ | 2 primitives are repeatedly applied: **rotational symmetry P** and **rotation±1**. No nesting. |
| 4SEGMENTS | 7 | $[[+0, H]^4\{-1\}]$ | The first segment is built from **axial symmetry**. It is translated four times shifting its starting point. Two embedded levels of regularity. |
| 4DIAGONALS | 7 | $[[+0, P]^4\langle +1 \rangle]^*$ | **Rotational symmetry P** is repeatedly applied to four consecutive starting points. Two embedded levels of regularity. |
| 2ARCS | 8 | $[[+0, [+1]^3]^2\langle B \rangle]^*$ | **Rotation±1** is applied 3 times. Global **axial symmetry** to determine the 4 remaining positions. Two embedded levels of regularity. |
| 2SQUARES | 8 | $[[+0, [+2]^3]^2\langle -1 \rangle]^*$ | **Rotation±2** is applied 3 times to draw the first square, then again after shifting the starting point by **rotation±1** to draw the second square. Two embedded levels of regularity. |
| 2RECTANGLES | 10 | $[[[+0, V]^2\langle P \rangle]^2\{-2\}]^*$ | A **vertical symmetry V** draws long segment, then a **rotational symmetry P** is applied in order to trace a first rectangle. Then the same sequence is replicated after a global **rotation±2** to trace the second rectangle. 3 embedded levels of regularity. |
| IRREGULAR | 16 | $[+0, P, +3, +2, P, A, V, H]$ | Mere list of the successive transitions between locations. No nesting. |

The table presents one of the descriptions (*) or the unique description provided by the language of geometry for all the sequences we considered in the study. A verbal description of the sequence structure is also provided.

### Assessing a possible contribution of eye movements to spatial decoding

During the MEG acquisition, participants were instructed to fixate the central cross while the items were flashed. Their gaze was monitored online to make sure that they did. However, involuntary eye movements may contaminate MEG recordings and lead to decodable nonbrain signals (Mostert et al., 2018; Quax et al., 2019). We assessed if the spatial decoding and the anticipation results were partly due to participants eye movements during the sequence presentation. We conducted control analyses on the participants for whom we could collect eye tracking data (14 out of 20).

We first measured gaze position during the sequence trials to assess its distance to the central fixation cross. Figure S1B shows the gaze heatmap overlaid onto the fixation cross and the 8 octagon vertices presented to the participants. For each participant separately, we computed the standard deviation of the distribution of distances to the fixation cross. The mean standard deviation was 38 pixels (SD = 23 pixels), which was larger than the fixation cross (30 pixels) but much smaller than the octagon radius (225 pixels). This confirmed that participants did not saccade to the sequence items.

Moreover, the performance of a position decoder trained on the eye tracking and EOG data was much lower than the one of a decoder trained on the MEG data (figure S1A and S1D). We conclude that, even if we cannot fully exclude that the late positive performance (from 220ms on) may correspond to eye movements and micro-saccades, they do not play a dominant role for the brain decoding results.

Finally, we ran a mutual information analysis to determine how much information was shared between the MEG decoding results and eye movement decoding results (figure S1C). The mutual information was calculated over the correctly classified MEG trials, therefore providing a measure of the amount of information provided by the eye movement decoding results about the MEG decoding results (Quax et al., 2019). Remarkably, the mutual information was lower in the time interval corresponding to the maximal performance of the MEG data decoder, suggesting that during that time window the MEG data decoder based its predictions on brain rather than eye signals.

**Figure S1:**
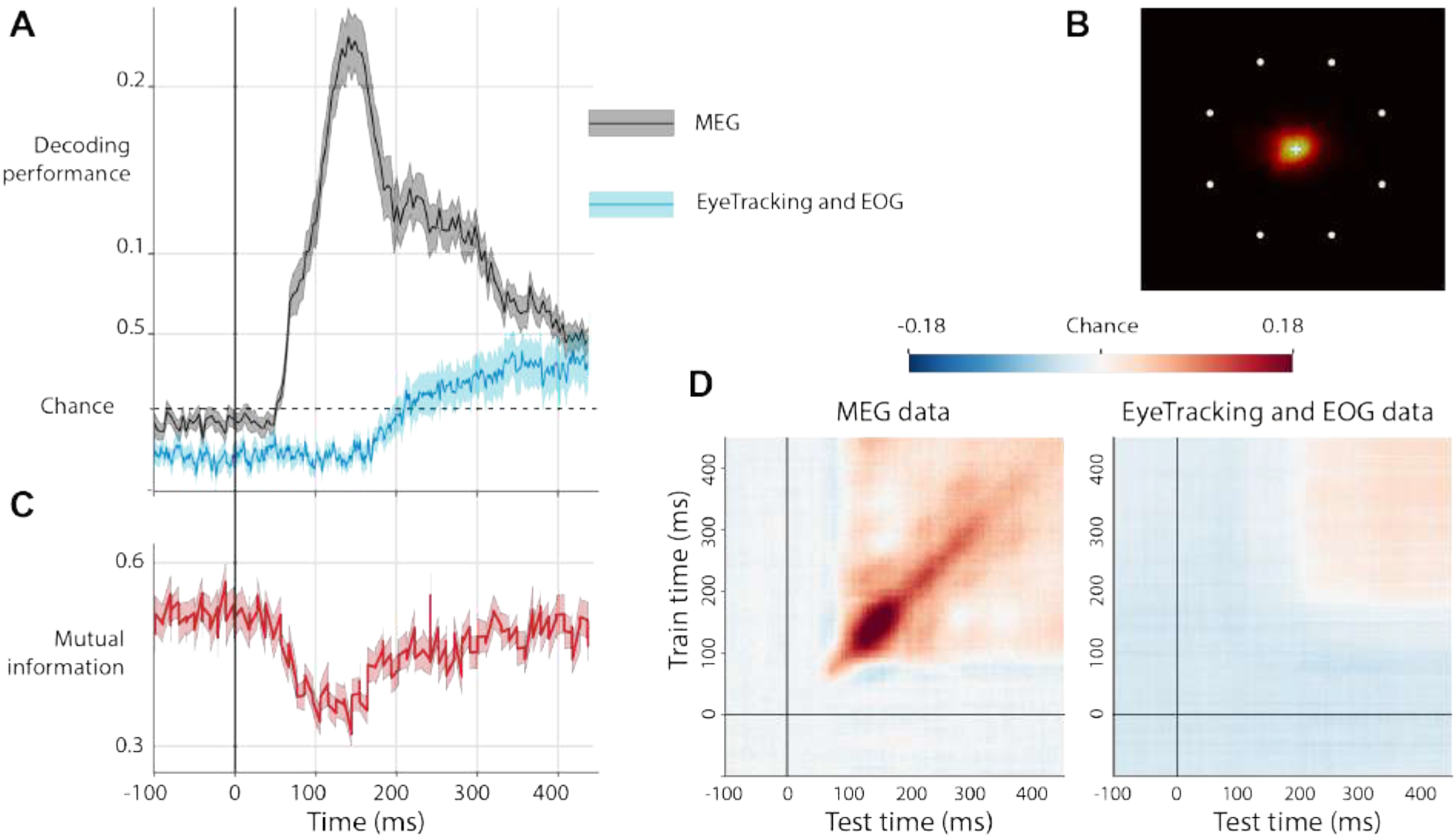
Eye movements do not account for brain decoding of stimulus location. A. Performance in decoding stimulus location from MEG data (black) and eye tracking and EOG data (blue) as a function of time following the flash of a dot at a given location. Decoding from MEG data was much better and significantly positive much earlier than decoding from eye tracking and EOG data. B. The gaze heatmap during the sequence blocks confirms that participants did not saccade to the sequence items. Instead, in agreement with the instructions, they maintained the gaze on the central fixation cross. C. When the MEG decoder had maximal performance, the mutual information, which quantifies the amount of information provided by the eye movements about the MEG decoding results, was reduced. This confirms that during that time window, the MEG data decoder based its predictions on brain rather than eye data. D. Average generalization across time matrix showing the location decoding score for MEG data (left) and eye tracking and EOG data (right) as a function of training times (y axis) and testing times (x axis). The score for MEG data is much larger and significantly positive much earlier than the one for eye tracking and EOG data.

### Controlling for possible confounding factors for the ordinal position decoding

As a control for Figure 7, we tested whether the ordinal decoding could be due to the temporal structure in any sequences, perhaps due to an excessive power of the decoders to detect signals. Thus, we labelled the repeat+1 and irregular items with pseudo-ordinal positions that were yoked to the ordinal positions, either of the 4segments and 4diagonals sequences (ordinal positions 1-2-1-2…) or of the 2arcs and 2squares sequences (ordinal positions 1-2-3-4…). No significant oscillations were detected when trying to decode the ordinal positions 1-2-1-2… (figure S4 panels A and B) but a peak at f/4 (p<0.001) and f/2 (p<0.05) could be found when decoding ordinal positions 1-2-3-4-1… suggesting that participants may have segmented these sequences into group of 2 or 4 in order to memorize them. To determine if these signals originated from the repeat+1 or the irregular sequence, we ran a separate analysis for each of them (figure S5). The power spectrum for the irregular sequence exhibited no peak at f/4 or f/2, as expected, but a very small but significant peak at f/2 was found for repeat+1, suggesting that this sequence, although monotonous, was perhaps parsed according to the sides of the regular octagon.

**Figure S2:**
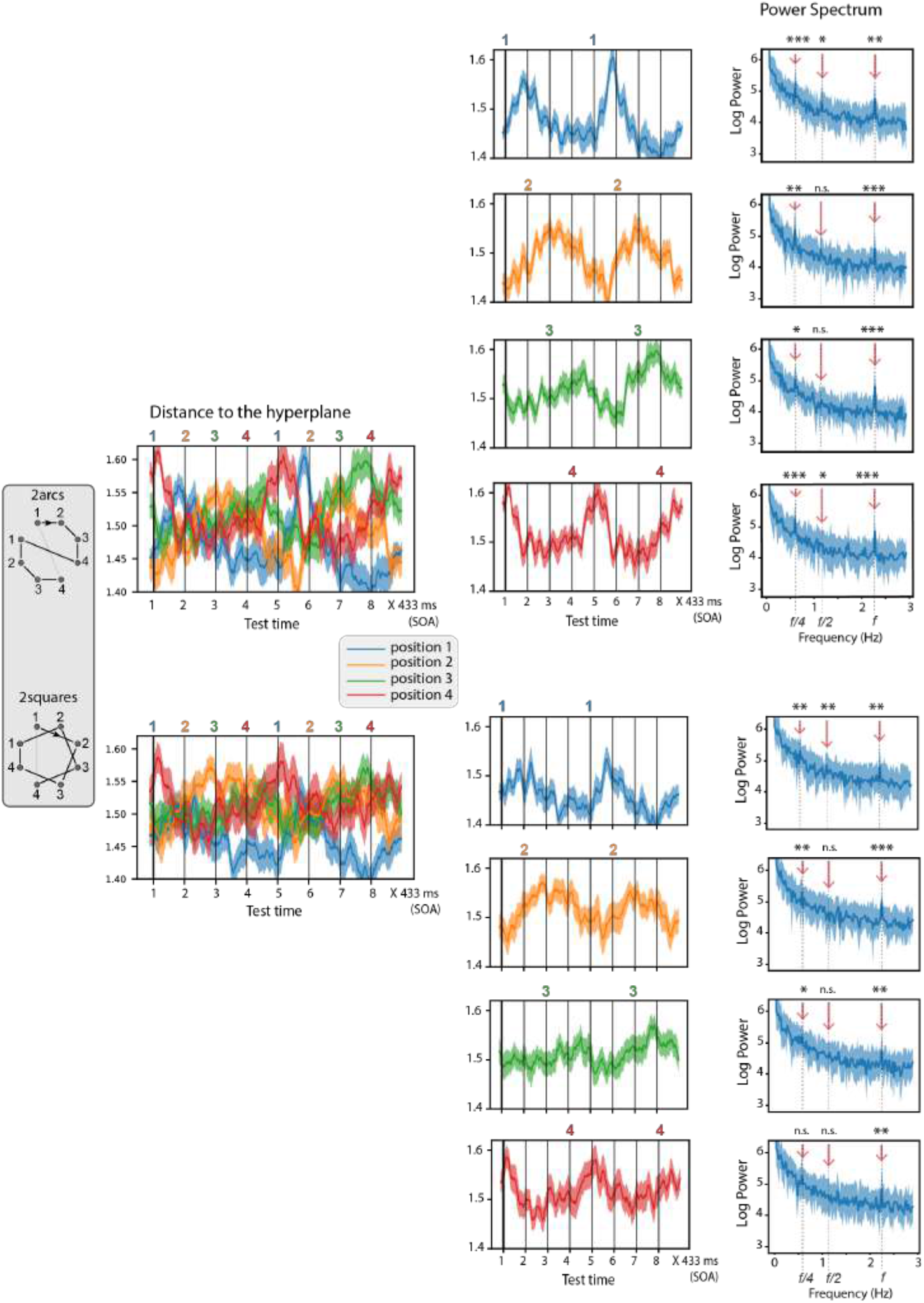
The ordinal code is reactivated in a periodic manner for each position. Decoders were trained to predict the ordinal positions 1-2-3 and 4 in the inner groups of sequences 2arcs and 2squares. The top figures were obtained by cross-validating across blocks, the bottom ones by generalizing across sequences. Left and middle columns show the time course of the average output of ordinal decoders trained in a time window 300-500 ms, during a sequence of 8 consecutive items. Blue, orange, green and red lines indicate 1^st^, 2^nd^, 3^rd^ and 4^th^ positions. The y axis shows the distance to the ordinal classification hyperplane for each decoder. The onsets of the sequence items are indicated by vertical lines, and times are reported from the onset of the first item (thick vertical line). Right column, power spectrum obtained from the Fourier transform of those time courses throughout the full presentation blocks for the two sequences. Statistics are provided at frequencies f = 2.31 Hz, f/2=1.15 Hz and f/4 = 0.58 Hz. Peaks at f/4 are present for the 2^nd^ and 3^rd^ ordinal positions, confirming that the effects observed in figure 7 do not only originate exclusively from the items that open and close a component.

**Figure S3:**
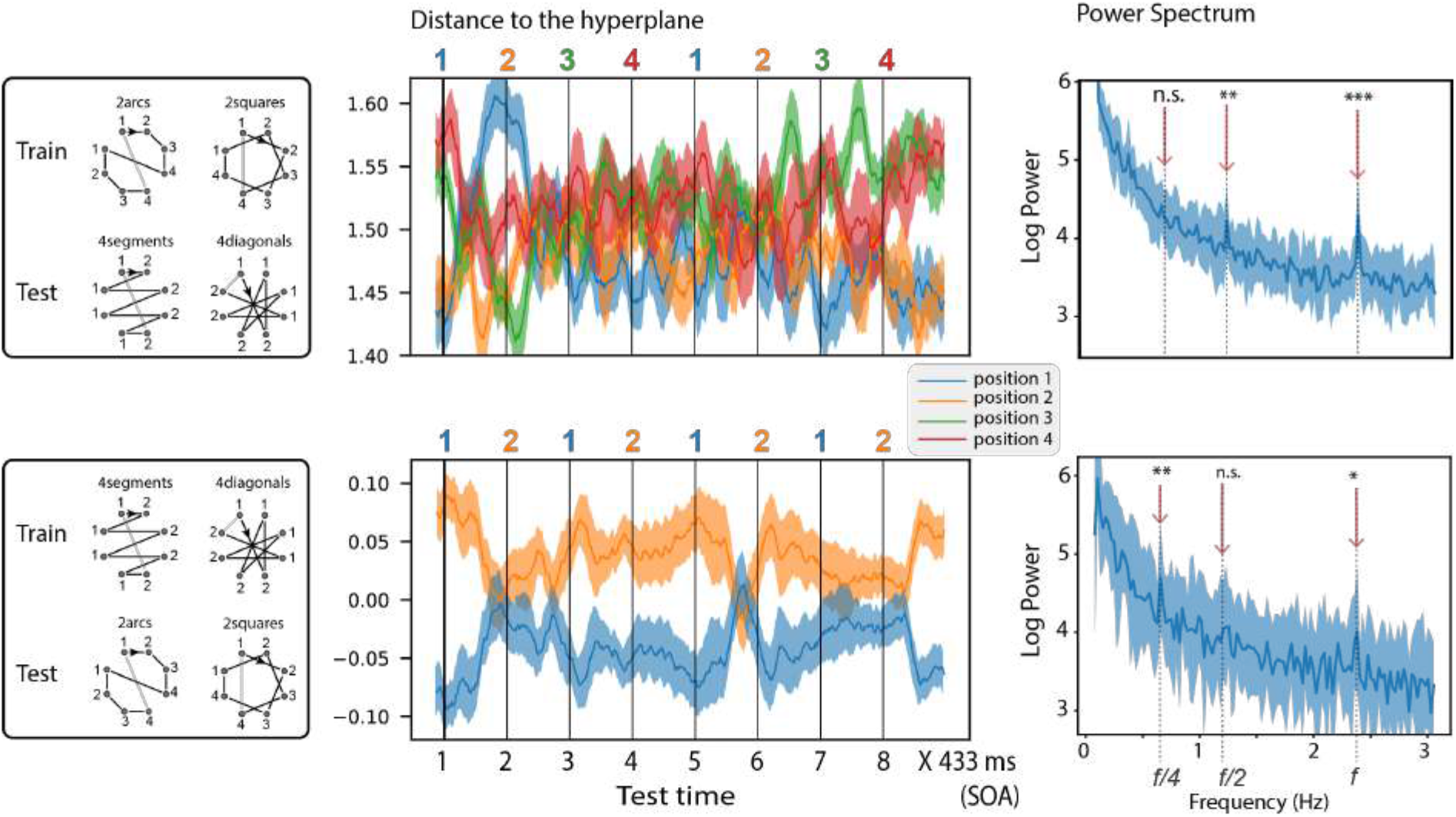
Generalization of the ordinal code across component size. Top panels, decoders were trained on 2arcs and 2squares sequences to predict the ordinal positions 1, 2, 3 and 4, and were then tested on 4segments and 4diagonals. Bottom panels, decoders were trained on 4segments and 4diagonals sequences to predict the ordinal positions 1 and 2, and were then tested on 2arcs and 2squares. The left column shows the time course of the average output of ordinal decoders trained in a time window 300-500 ms, during a sequence of 8 consecutive items. Blue, orange, green and red lines indicate the outputs of decoders for the 1^st^, 2^nd^, 3^rd^ and 4^th^ positions. The y axis shows the distance to the ordinal classification hyperplane for each decoder. The onsets of the sequence items are indicated by vertical lines, and times are reported from the onset of the first item (thick vertical line). The top left panel shows that codes for the 1^st^ and 4^th^ positions obtained from 2arcs and 2squares are active for the first and the last items of the 4segments and 4diagonals sequences. The top right panel shows the power spectrum obtained from the Fourier transform of the distance time-series. A significant peak is present at f/2 and corresponds to the reactivation of the codes for the 1^st^, 2^nd^ and 4^th^ positions (p<0.01 at f/2 for these 3 positions). The bottom left panel shows that codes for the 1^st^ and 2^nd^ positions obtained from 4segments and 4diagonals sequences reactivate for the same positions of 2arcs and 2squares sequences (with the code for 2^nd^ position showing a temporally extended response, sustained from positions 2 to 4 in the generalization sequences). This finding is confirmed by the presence of a peak at f/4 in the power spectrum of the distance time-series.

**Figure S4:**
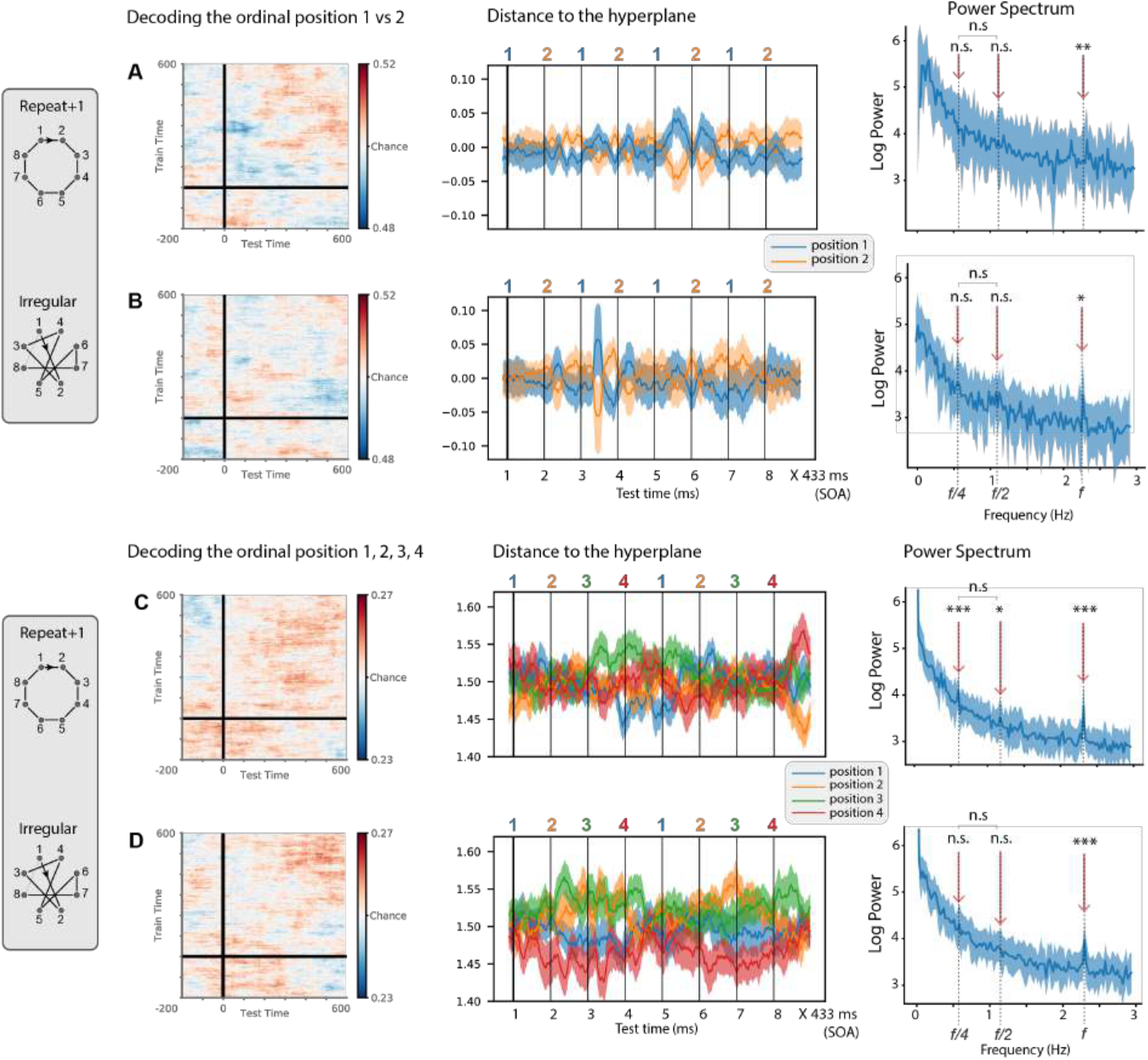
Ordinal position code, control analysis. In this control analysis, with the same format as figure 7, the same decoders that were used to predict actual ordinal position in regular sequences (4 segments, 4 diagonals, 2arcs and 2squares) were now trained with data that, according to our model, should not contain ordinally organized subgroups, i.e. MEG data from the sequences repeat+1 and irregular. Panels A and C were obtained by cross-validating across blocks and B and D by generalizing across sequences. Format is similar to figure 7. Note the lack of significant decoding, or of significant peaks in the spectrograms, except for very small peaks but significant peaks at f/2 and f/4 in panel C. The further analysis presented in the next figure (S5) shows that the latter may arise from the repeat+1 sequences, which may have been parsed into groups of 2 or 4, perhaps according to left versus right or top versus bottom, thus separating the four sides of the octagon.

**Figure S5:**
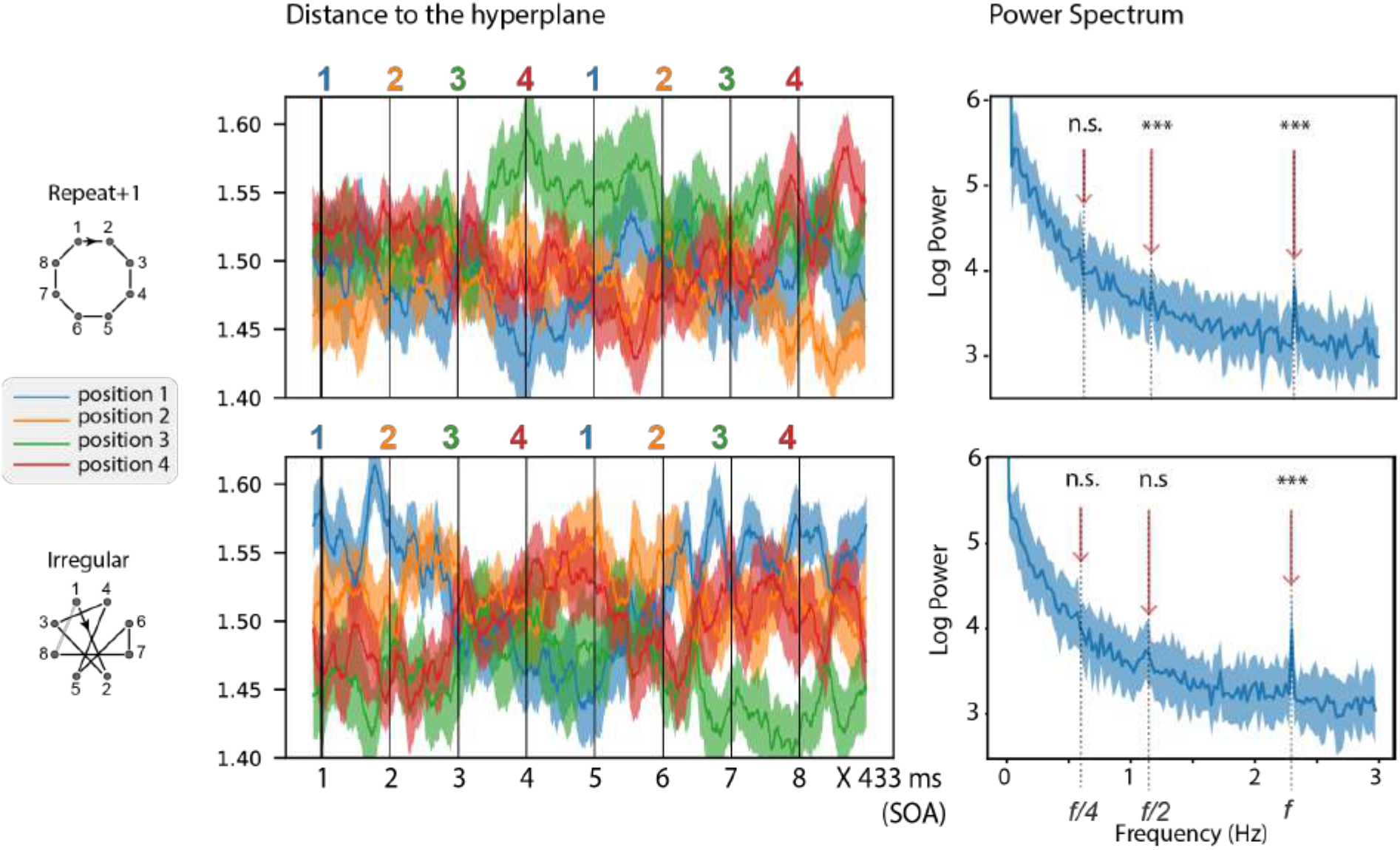
Ordinal position code 1-2-3-4, control analysis separately for repeat+1 and irregular sequences. In this control analysis, decoders were trained to predict the ordinal positions 1-2-3-4 separately for repeat+1 (top panels) and irregular (bottom panels) sequences. Note that the amount of training data is therefore halved, compared to previous analyses, and therefore the results are noisier and must be taken with caution. The left column shows the time course of the average output of ordinal decoders trained in a time window 300-500 ms, during a sequence of 8 consecutive items. Blue, orange, green and red lines indicate 1^st^, 2^nd^, 3^rd^ and 4^th^ positions. The y axis shows the distance to the ordinal classification hyperplane for each decoder (same format as figures 7 and S4). Although no pattern is easily discernable, the spectral analysis (right) detects a very small but significant peak at f/2 in repeat+1 sequences, which may correspond to a grouping by 2 of the sides of the octagon.

